# Cellular Dynamics and Genomic Identity of Centromeres in the Cereal Blast Fungus

**DOI:** 10.1101/475574

**Authors:** Vikas Yadav, Fan Yang, Md. Hashim Reza, Sanzhen Liu, Barbara Valent, Kaustuv Sanyal, Naweed I. Naqvi

## Abstract

A series of well-synchronized events mediated by kinetochore-microtubule interactions ensure faithful chromosome segregation in eukaryotes. Centromeres scaffold kinetochore assembly and are among the fastest evolving chromosomal loci in terms of the DNA sequence, length, and organization of intrinsic elements. Neither the centromere structure nor the kinetochore dynamics is well studied in plant pathogenic fungi. Here, we sought to understand the process of chromosome segregation in the rice blast fungus, *Magnaporthe oryzae*. High-resolution confocal imaging of GFP-tagged inner kinetochore proteins, CenpA and CenpC, revealed an unusual albeit transient declustering of centromeres just before anaphase separation in *M. oryzae*. Strikingly, the declustered centromeres positioned randomly at the spindle midzone without an apparent metaphase plate *per se*. Using chromatin immunoprecipitation followed by deep sequencing, all seven centromeres were identified as CenpA-rich regions in the wild-type Guy11 strain of *M. oryzae*. The centromeres in *M. oryzae* are regional and span 57 to 109 kb transcriptionally poor regions. No centromere-specific DNA sequence motif or repetitive elements could be identified in these regions suggesting an epigenetic specification of centromere function in *M. oryzae*. Highly AT-rich and heavily methylated DNA sequences were the only common defining features of all the centromeres in rice blast fungus. PacBio genome assemblies and synteny analyses facilitated comparison of the centromere regions in distinct isolate(s) of rice blast, wheat blast, and in *M. poae.* Overall, this study identified unusual centromere dynamics and precisely mapped the centromere DNA sequences in the top model fungal pathogens that belong to the *Magnaporthales* and cause severe losses to global production of food crops and turf grasses.

**Author summary:** *Magnaporthe oryzae* is an important fungal pathogen that causes an annual loss of 10-30% of the rice crop due to the devastating blast disease. In most organisms, kinetochores are arranged either in the metaphase plate or are clustered together to facilitate synchronized anaphase separation of chromosomes. In this study, we show that the initially clustered kinetochores separate and position randomly prior to anaphase in *M. oryzae*. Centromeres, identified as the site of kinetochore assembly, are regional type without any shared sequence motifs in *M. oryzae*. Together, this study reveals atypical kinetochore dynamics and identifies functional centromeres in *M. oryzae*, thus paving the way to define heterochromatin boundaries and understand the process of kinetochore assembly on epigenetically specified centromere loci in the economically important cereal blast and summer patch pathogens. This study paves the way for understanding the contribution of heterochromatin in genome stability and virulence of the blast fungus.

## Introduction

Faithful chromosome segregation is one of the essential processes required for maintaining genome integrity in dividing cells. This process is successfully carried out by the attachment of microtubules, emanating from opposite spindle poles, to the proteinaceous multi-subunit structure, the kinetochore, that is pre-assembled onto centromeres [1, 2]. The centromere forms a crucial part of this machinery and yet, it is one of the most rapidly evolving loci in eukaryotic genomes [3, 4]. On the contrary, the proteins that bind to centromere DNA are evolutionary conserved [2]. Centromere DNA shows a wide diversity in the length and composition of the underlying DNA sequence. A few fungal species, like *Saccharomyces cerevisiae*, harbor centromeres that are less than 400 bp comprising of conserved DNA sequence elements to form point centromeres [5]. Most others possess regional centromeres that span from a few kilobases to several megabases. Unlike point centromeres, regional centromeres in an organism can span hundred of kbs and are often are defined by epigenetic factors. For example, the regional centromeres in *Schizosaccharomyces pombe* and *Candida tropicalis* have a homogenized central core flanked by inverted repeats [6, 7]. Likewise, the regional centromeres in *Cryptococcus neoformans* possess specific retrotransposons that are present randomly therein [8]. In contrast, centromeres in *Candida albicans*, *Candida lusitaniae,* and *Candida dubliniensis* differ between all chromosomes and lack a conserved DNA sequence element [9-11]. Centromeres in filamentous fungi like *Neurospora crassa*, on the other hand, span long stretches of repetitive DNA but lack a consensus sequence or pattern [12, 13]. Metazoans and plants also have regional centromeres that are up to few megabases long, and mostly consist of repetitive DNA or transposons [14-16].

Despite this sequence divergence, centromeres in most studied organisms are bound by a centromere-specific histone H3 variant CENP-A /CenH3/Cse4, also known as the hallmark of centromere identity [4, 17]. CENP-A forms the foundation of the kinetochore assembly and is essential for cell viability in all organisms studied until date. Evolutionary conservation of CENP-A along with other kinetochore proteins also provides an efficient tool to identify centromeres. Additionally, studies with fluorescently-labeled inner kinetochore proteins such as CENP-A or CENP-C/Cen-C/Mif2 has led to an understanding of spatial dynamics of the kinetochore within the nucleus [18-22]. These studies established that kinetochores in most yeast species are clustered throughout the nuclear division, and unlike metazoan *CEN*, do not align on a metaphase plate. However, more recently, some variations to the metaphase plate or kinetochore clustering have been reported revealing the diversity in this phenomenon. Kinetochores remain clustered throughout the cell cycle in two well-studied ascomycetes, *S. cerevisiae,* and *C. albicans* [23, 24]. In *S. pombe,* kinetochores undergo a brief declustering during mitosis but remain clustered otherwise [18, 25]. Another ascomycete, *Zymoseptoria tritici*, shows multiple kinetochore foci instead of a single cluster during interphase although their localization dynamics during mitosis remains unexplored [26]. On the other hand, the cells of a basidiomycete *C. neoformans* display multiple foci of kinetochores in interphase, but kinetochores gradually cluster during mitosis [19, 22]. Even the phenomenon of centromere/kinetochore clustering is observed in *Drosophila* that depends on centric chromatin rather than specific DNA sequences [27].

Besides CENP-A, several other chromatin features are known to be associated with centromeres. For example, centromeres are devoid of genes/ORFs and exhibit a significantly low level of polyA transcription as compared to the rest of the genome [8, 28]. Furthermore, centromeres in many organisms are heterochromatic in nature and harbor the heterochromatic marks like H3K9di/trimethylation and DNA methylation [8, 13, 29]. A preference for AT-rich DNA sequence is evident for centromere formation in some organisms [13, 30-32]. It is important to note that none of these features exclusively define centromeres and, in most cases, the importance of an individual factor in defining centromere loci is not well understood. However, the presence of such features on discrete chromosomal loci may pave the way for predicting centromeres in organisms in which genome tractability is difficult.

Magnaporthales is an order of ascomycete fungi comprising of many important plant pathogenic species including *Magnaporthe oryzae* (synonym of *Pyricularia oryzae*) and *Magnaporthe poae* (synonym of *Magnaporthiopsis poae*) [33]*. M. oryzae* includes host-adapted lineages (pathotypes) causing the devastating blast diseases in cereal crops including rice, wheat, barley and millets [34-36]. *M. poae* is responsible for summer-patch disease in turf grasses [37]. The *M. oryzae* lineage causes rice blast which remains a constant threat to agriculture-based economies due to significant damage to rice harvests. Recently, wheat blast disease, caused by the *M. oryzae Triticum* lineage has emerged as a major threat to global wheat production [38]. Rice blast has also become a model pathosystem for studying host-pathogen interactions due to the availability of the genome sequence, fully characterized infection cycle, genetic tractability and economic significance of the fungus [39, 40]. However, even with the availability of the genome sequence and annotated assembly, the centromere/kinetochore identity of the blast fungus remains unexplored or poorly defined. Here, we first studied and characterized orthologs of CENP-A and CENP-C, two well-conserved kinetochore proteins, to understand the kinetochore dynamics in this organism and used these kinetochore proteins as tools to identify bona fide centromeres.

## Results

### Kinetochores are clustered during interphase in *M. oryzae*

A subset of putative kinetochore proteins was previously annotated in *M. oryzae* [12]. We expanded the list further by identifying putative orthologs of the additional conserved kinetochore proteins using *in silico* predictions (S1 Table). Multiple sequence alignment established the identity of at least two most conserved inner kinetochore proteins: CenpA (MGG_06445, a homolog of CENP-A) and CenpC (MGG_06960, a homolog of CENP-C) (S1 Fig) in *M. oryzae*. CenpA and CenpC of *M. oryzae* share 73% and 42% sequence identity with their *N. crassa* counterparts CenH3 and CEN-C, respectively. Next, we functionally expressed the GFP-tagged CenpA and CenpC from their native genomic loci in the wild-type Guy11 strain of *M. oryzae*. GFP-CenpA and CenpC-GFP signals appeared as single dot-like and co-localized on chromatin, marked by mCherry-tagged histone H1 (Fig 1A and 1B). Further, co-localization of CenpA and CenpC signals confirmed their overlapping spatial positions in both mycelia and conidia (Fig 1C). Clustering of kinetochores is a hallmark feature of many yeast and fungal genera. Such clustered kinetochores are often found in close proximity to the spindle pole bodies (SPBs) [41]. We localized SPBs by tagging Alp6 (MGG_01815, an ortholog of *S. cerevisiae* Spc98) with mCherry and observed that SPBs localize close to the clustered GFP-CenpA signals in *M. oryzae* (Fig 1D). These results indicate that kinetochore localization during interphase in *M. oryzae* is similar to that observed in other ascomycetes. Our attempts to delete *CENPA* or *CENPC* in *M. oryzae* failed indicating that both are essential for cell viability. This result was further corroborated by conditional repression of *CENPA* using the Tet-off system. The *Tet-GFP-CENPA* strain ceased to grow on culture media supplemented with doxycycline, the condition in which Tet-driven *CENPA* expression is shut down (S2A Fig). Overall, the conserved sequence features and the subcellular localization patterns confirmed that CenpA and CenpC are evolutionarily conserved kinetochore proteins in *M. oryzae*.

**Fig 1.**
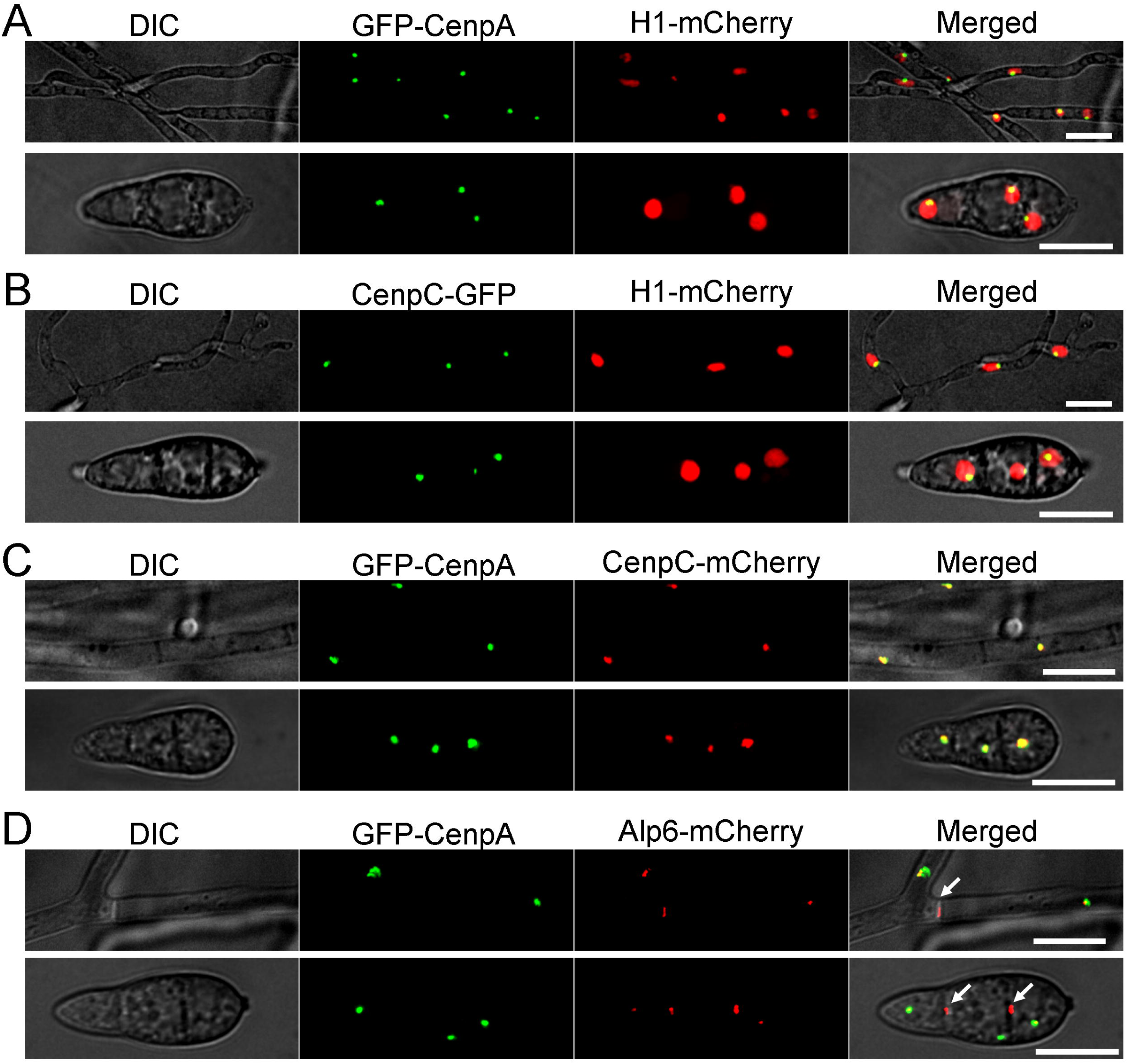
Localization patterns of CenpA and CenpC reveal kinetochores are closely associated with each other in *M. oryzae*. **(A)** The *M. oryzae* strain MGYF03 exhibits a single dot-like GFP-CenpA signal localized at the periphery of each nucleus marked by mCherry-histone H1 in both mycelia (upper panel) and conidia (lower panel). **(B)** Similarly, another inner kinetochore protein CenpC-GFP in the strain MGYF04 is found to be localized at the periphery of the mCherry-histone H1 marked nucleus in both mycelia (upper panels) and conidia (lower panels). **(C)** Co-localization of GFP-CenpA and CenpC-mCherry revealed complete overlapping signals in both mycelia and conidia in the MGYF05 strain. **(D)** In the strain MGYF08, the clusters of GFP-CenpA are closely associated with the spindle pole body (SPB) component Alp6-mCherry. In addition to SPBs, the Alp6 signals were also observed at the septa (white arrows). The fluorescence images shown here are maximum projections from Z stacks consisting of 0.5 µm-spaced planes. Scale bar = 10 μm.

### Kinetochores in *M. oryzae* undergo declustering-clustering dynamics during mitosis

To study the cellular dynamics of kinetochores in *M. oryzae*, we localized microtubules by expressing mCherry-TubA or GFP-TubA fusion protein and co-localized it with GFP-CenpA. During interphase, the microtubules are mostly localized throughout the cytoplasm (S2B Fig). Live-cell imaging during mitosis revealed dispersed GFP-CenpA signals localized along the mitotic spindle (Fig 2A and S1 Movie). Strikingly, the declustered dot-like signals of GFP-CenpA then segregated into two halves in a non-synchronous manner. Once segregated, the signals began to cluster again and localized as two bright foci close to poles of the mitotic spindle. To further probe the dynamics of kinetochore segregation, we performed high-resolution imaging in mitotic cells expressing GFP-CenpA (Fig 2B, C and S2, S3 Movie). We observed that while the GFP-CenpA signals were spread out, they were localized in pairs, most likely representing the segregated kinetochore signals (Fig 2B, time 00:32). We were able to count fourteen discrete spots of GFP-CenpA corresponding to 14 kinetochores of the seven duplicated chromosomes. These results suggest that kinetochores in *M. oryzae* remain largely unclustered during mitosis. It was further supported by co-localization of GFP-CenpA with a SPB markerAlp6-mCherry during the mitotic stages (Fig 2D). In pre-mitotic cells, we observed two duplicated spots of Alp6-mCherry that co-localized with replicated clustered GFP-CenpA signals. During mitosis, GFP-CenpA signal localized as multiple puncta scattered in between the two SPBs represented by Alp6-mCherry. After the division, the GFP-CenpA/kinetochores clustered again and localized adjacent to the SPBs (S2C Fig and S4 Movie). Taken together, we conclude that kinetochores decluster during mitosis in *M. oryzae,* and align along the mitotic spindle. Furthermore, we infer that an equatorial plate alignment of the kinetochores is not evident in *M. oryzae*, indicating a lack of a well-defined metaphase plate. Similar dynamics of the kinetochore and microtubules were observed in *M. oryzae* cells during pathogenic development and *in planta* conditions (S3 Fig, S5 and S6 Movie). Based on these observations, we propose a schematic model for the kinetochore and SPB dynamics during the mitotic cycle in rice blast where the dynamics of kinetochore clustering-declustering is most likely dependent on their direct link to the SPBs (Fig 2E). During mitosis, this link is likely broken, and the clustering is thus perturbed. We infer that such timely and dynamic kinetochore clustering/declustering is crucial for proper chromosome segregation in *M. oryzae.*

**Fig 2.**
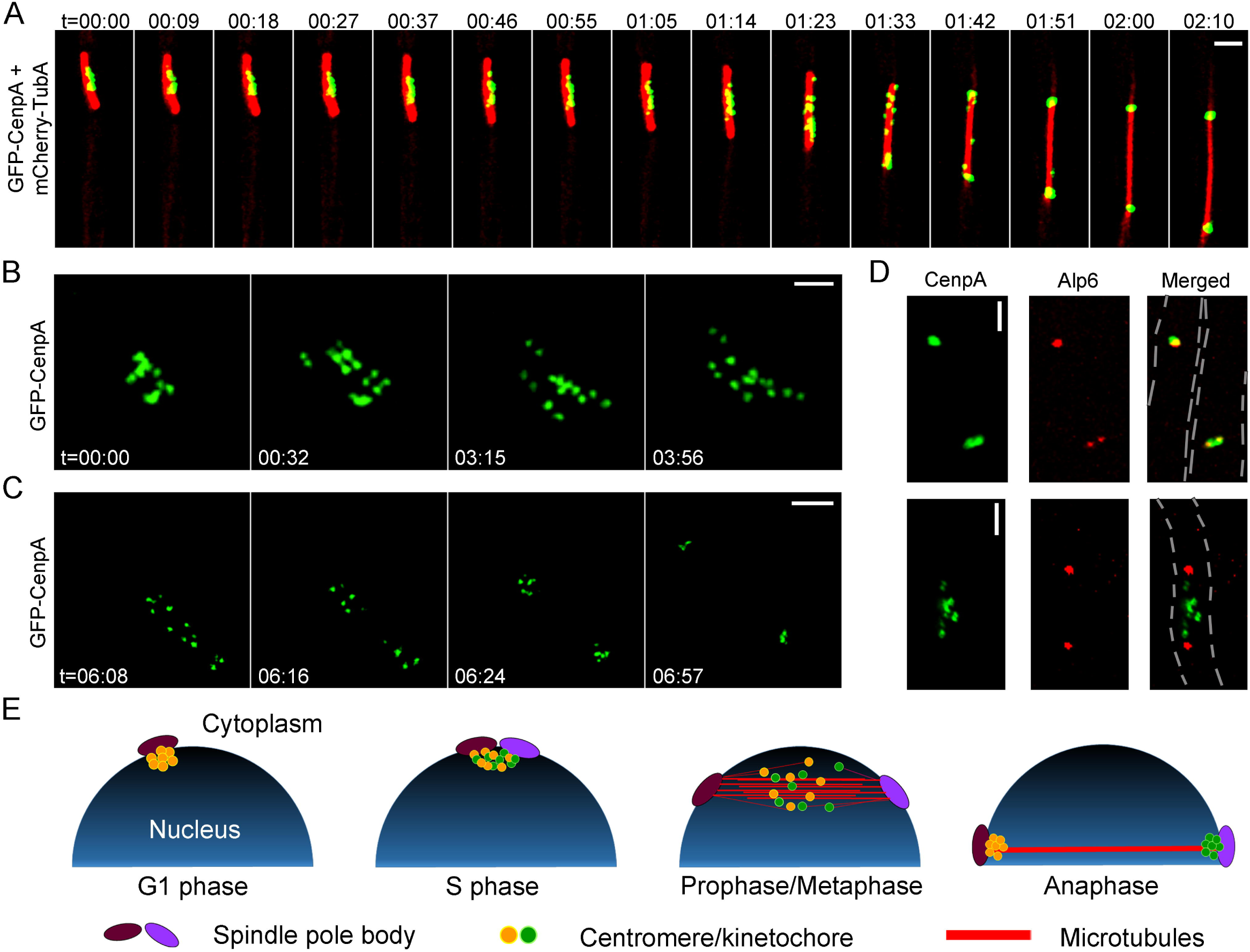
Kinetochores decluster momentarily but arrange randomly on the spindle axis before sister kinetochore separation during anaphase in *M. oryzae*. **(A)** Time-lapse imaging of strain MGYF07 cells exhibited that the GFP-CenpA signals separate from each other and move along the mitotic spindle (mCherry-TubA). Also see movie S1. The images shown are maximum projections of 0.3 µm-spaced *Z* stacks. t = minutes:seconds. Scale bar = 2 µm. **(B)** High-resolution time-lapse images showing the declustering of kinetochores (GFP-CenpA) during the process of mitosis in strain MGYF01 (Also, see Movie S2). The images were acquired with *Z* projections of 0.17-µm step size. t = minutes:seconds. Scale bar =1 µm. **(C)** High-resolution time-lapse images of MGYF01 cells showing the segregation dynamics of sister kinetochores in daughter cells during metaphase to anaphase transition and finally reclustering of kinetochores in post-anaphase cells (Also see movie S3). t = minutes:seconds. Scale bar = 2 µm. **(D)** Spatial organization of kinetochores (GFP-CenpA) and SPBs (Alp6-mCherry) in strain MGYF08 during pre-mitotic stage (upper panel) and early mitosis (lower panel). Scale bar = 2 µm. **(E)** A schematic depiction of centromere dynamics at specific stages of the cell cycle in *M. oryzae*. For simplification, chromosomes and astral microtubules are omitted in the schematic.

### Kinetochore protein binding identifies regional centromeres in *M. oryzae*

CenpA binding is a hallmark of functional centromeres in eukaryotes [4, 15]. We used GFP-CenpA as a tool for molecular identification of centromeres in the *M. oryzae* genome. We utilized chromatin immunoprecipitation (ChIP) assays followed by deep sequencing (ChIP-seq) of GFP-CenpA-associated chromatin fragments and aligned the reads on the recently published PacBio genome assembly of the wild-type Guy11 strain of *M. oryzae* [42]. This analysis revealed seven distinct CenpA-rich regions across the genome, one each on seven different contigs (Fig 3, Table 1 and S4 Fig). The CenpA binding spans a 57 to 109 kb region suggesting that *M. oryzae* possesses large regional centromeres. The centromere identity of these regions was further validated independently by binding of another evolutionarily conserved kinetochore protein CenpC. ChIP-qPCR using the fungal strain expressing CenpC-GFP (Fig 3C) confirmed specific overlapping binding of CenpA and CenpC on each of these seven *CEN* regions. We also noticed an extra albeit short region of 1200 bp on Contig 4 apart from the seven distinct peaks in CenpA ChIP-seq analysis. The enriched peak mapped to the gene encoding the vacuolar morphogenesis protein AvaB (MGG_01045). Using specific ChIP-qPCR primers for this region, the aforementioned CenpA enrichment on Contig 4 was confirmed to be an artifact (Fig 3D). Overall, the binding of two independent kinetochore proteins at seven long regions confirmed that these are indeed authentic centromeres of the corresponding chromosomes in *M. oryzae*.

**Table 1.** Length and %GC content of centromeres in distinct *Magnaporthe* isolates.

| Isolate/<br>species | Guy11 |  | 70-15* | FJ81278** |  | B71*** |  | *M. poae* **** |  |
| --- | --- | --- | --- | --- | --- | --- | --- | --- | --- |
| *CEN*# | Coordinates | %GC | Coordinates | Coordinates | %GC | Coordinates | %GC | Coordinates | %GC |
| *CEN1* | Contig15:<br>571735-678893<br>(1,07,159) | 29.2 | Chr1:<br>4669580-4690159<br>(20,580) | Contig6:<br>942880-1032698<br>(89,819) | 26.8 | Chr1:<br>5247711-5361150<br>(113,440) | 29.4 | GL876966:<br>1947276-2058771<br>(111,496) | 25.5 |
| *CEN2* | Contig7:<br>313767-411084<br>(97,318) | 30.8 | Chr2:<br>451419-471909<br>(20,491) | Contig1:<br>4838642-4937646<br>(99,005) | 30.5 | Chr2:<br>397914-483923<br>(86,010) | 27.4 | GL876967:<br>2010105-2095459<br>(85,355) | 24.6 |
| *CEN3* | Contig2:<br>3795849-3894639<br>(98,791) | 33.0 | Chr3:<br>5534011-5547746<br>(13,736) | Contig13:<br>469353-567903<br>(98,551) | 34.6 | Chr3:<br>6398799-6492490<br>(93,692) | 25.8 | GL876968:<br>291161-385721<br>(94,561) | 23.8 |
| *CEN4* | Contig10:<br>1000090-1063263<br>(63,174) | 30.5 | Chr4:<br>845436-854002<br>(8,567) | Contig14: 1-38471 +<br>Contig3: 1-30023<br>(68,694) | 30.0 | Chr4:<br>815420-895650<br>(80,231) | 24.8 | GL876971:<br>2575155-2652612<br>(77,458) | 24.6 |
| *CEN5* | Contig4:<br>4014470-4071774<br>(57,305) | 28.0 | Chr5:<br>296487-302043<br>(5,557) | Contig4:<br>259241-318616<br>(59,376) | 28.8 | Chr5:<br>296404-367652<br>(71,249) | 24.8 | GL876972:<br>961971-1029283<br>(67,313) | 23.3 |
| *CEN6* | Contig5:<br>343714-452391<br>(1,08,678) | 32.3 | Chr6:<br>3852307-3861083<br>(8,777) | Contig12:<br>708384-794489<br>(86,106) | 29.3 | Chr6:<br>5687241-5758152<br>(70,912) | 24.8 | GL876975:<br>266207-359379<br>(93,173) | 24.2 |
| *CEN7* | Contig13:<br>573351-645424<br>(72,074) | 30.5 | Chr7:<br>2771164-2774628<br>(3,465) | - |  | Chr7:<br>3255069-3336736<br>(81,668) | 26.2 | GL876978:<br>1-33204<br>(33,204) | 37.1 |
|  |  |  |  |  |  | Scaffold 1:<br>40795-109519<br>(68,724) |  | GL876979:<br>1-17025<br>(17,025) | 24.6 |

Centromere numbers are according to the 70-15 genome assembly.
Numbers in the bracket represent centromere length in base pair.
\*Centromeres in 70-15 isolate contain breaks and hence %GC is not calculated for its centromeres.
\*\**CEN7* in FJ81278 could not be identified due to poor genome assembly.
\*\*\*Predicted centromere for mini-chromosome of B71 is also listed.
\*\*\*\*Two of the putative centromeres in *M. poae* are present at the end of two separate contigs.
Both are listed as different centromeres here.

**Fig 3.**
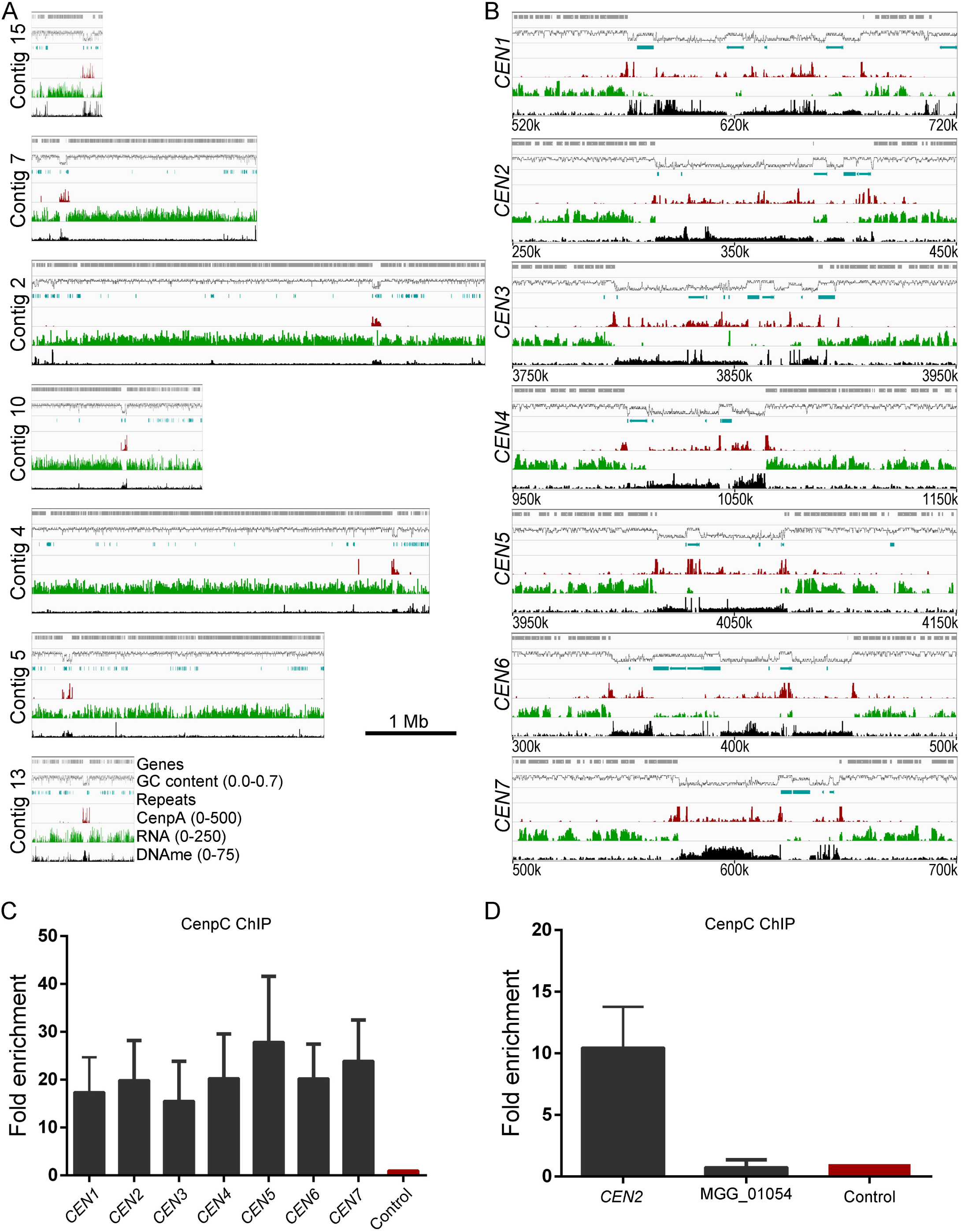
Identification of centromeres in *M. oryzae*. **(A)** Reads obtained from the GFP-CenpA ChIP-sequencing in the cross-linked mycelia of the strain MGYF01 identified one distinct enriched region on each of the seven contigs when aligned to the Guy11 genome assembly (see Fig S4 for the remaining contigs). CenpA-bound regions overlap with AT-rich, poorly-transcribed regions on each contig, and harbor 5mC DNA methylation (see text for details). The numbers in the bracket with parameters represent the minimum and maximum value along the *y*-axis. **(B)** The zoomed view of centromere regions in Guy11 depicting the presence of repeat elements, CenpA enrichment, polyA transcription and DNA methylation (5mC) status in these regions. A 200 kb region spanning the centromere is shown for each chromosome. The only common defining sequence feature of centromeres is AT-richness. **(C)** CenpC ChIP-qPCR analysis of cross-linked mycelia of the strain MGYF02 confirmed the centromere identity of each of the seven chromosomes of Guy11. Each bar represents the extent of enrichment obtained by one primer pair amplifying a unique sequence of each CenpA-bound region identified from the ChIP-seq analysis, and fold-enrichment values are normalized using a non-centromere region (ORF no. MGG_01917) as a control. Error bars represent the standard deviation calculated from three independent experiments. **(D)** ChIP-qPCR results showed that the gene MGG_01045 is not enriched with CenpC as observed in CenpA ChIP-seq analysis.

A detailed analysis revealed that the seven centromeres in *M. oryzae* comprise of highly AT-rich sequences (≥67%) (Fig 3B and Table 1). The centromeres in *M. oryzae* harbor a few repetitive elements (Fig 3B, S1 Dataset). However, these elements are neither exclusive to the centromeres nor common among the seven centromeres in *M. oryzae*. Further in-depth analysis of these regions did not reveal any common DNA sequence motif or repeats as supported by the dot-plot analysis of each centromere (S5 Fig). We then examined the transcriptional status and base modifications associated with centromeric chromatin using the published RNA-sequencing and bisulfite sequencing data [43, 44]. Centromeres in *M. oryzae* are found to be poorly transcribed and harbor 5mC DNA methylation (Fig 3). Based on these results, we conclude that centromeres in *M. oryzae* do not share any common DNA sequence motif or repeat element and that AT-richness is likely the only defining sequence feature of all the centromeres in *M. oryzae*. Additionally, we also infer that centromeres in *M. oryzae* are large, regional and lie within transcriptionally-poor 5mC-rich DNA regions of the genome.

### Centromere DNA sequences are rapidly evolving in rice blast isolates

The MG8 genome assembly is based on the sequencing of the *M. oryzae* isolate 70-15, which represents a progeny of the Guy11 strain [39, 45, 46]. The PacBio genome sequence of Guy11 provides a near complete end-to-end chromosome-wide coverage of the 70-15 genome, the only chromosome-level sequence assembly (MG8) available for an *M. oryzae* rice pathogen (S6A Fig). Thus, we attempted to identify the centromere location in the 70-15 isolate by aligning CenpA ChIP-seq reads on to the MG8 assembly. This analysis revealed seven distinct peaks, one on each chromosome (Fig 4 and Table 1). We also observed two additional CenpA-enriched regions in the unassembled Supercontig8.8 of MG8 assembly for 70-15 (S6B Fig). Additionally, the identified centromere on chromosome 7 in this assembly matched the region previously predicted to harbor the centromere based on genetic analysis [47].

**Fig 4.**
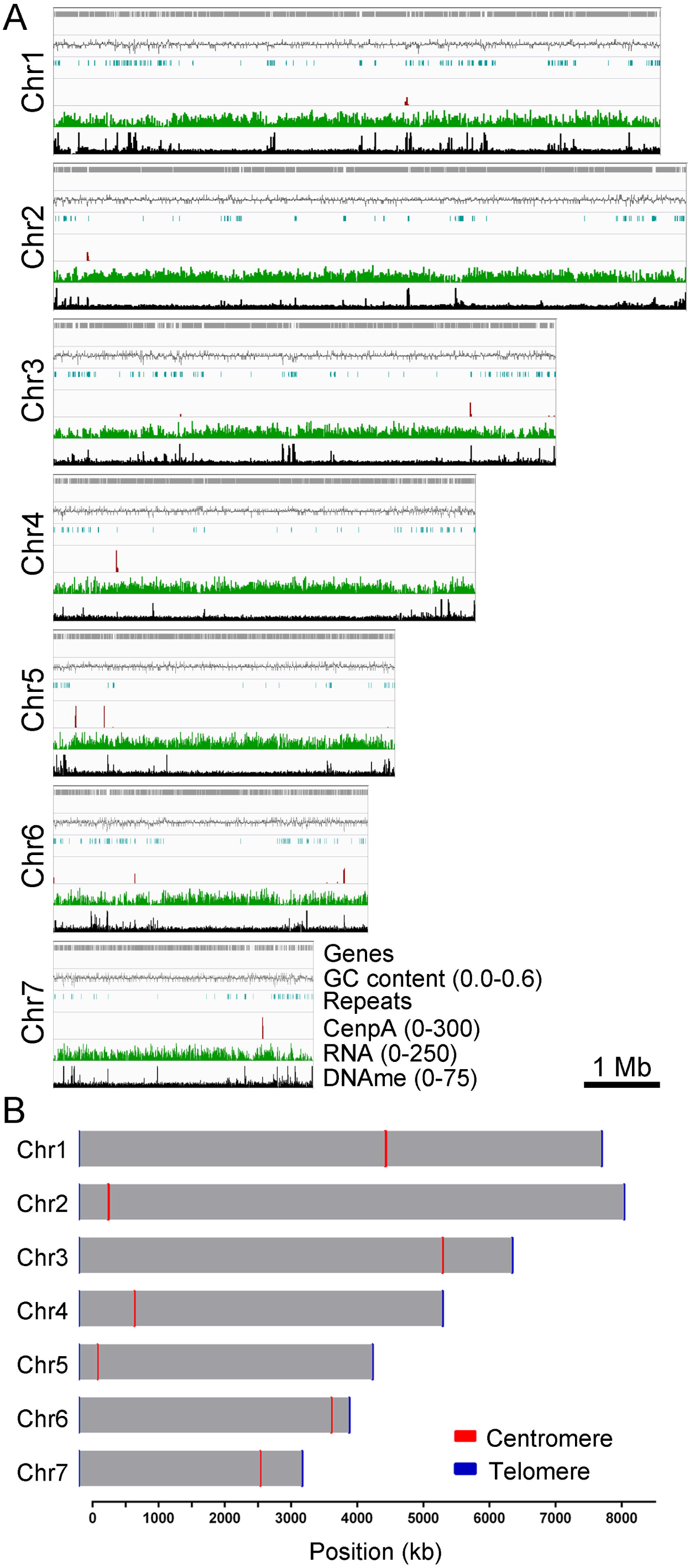
Identification of centromeres in *M. oryzae* strain 70-15 (MG8 assembly; Broad Institute). **(A)** Mapping of GFP-CenpA ChIP-seq reads to the reference MG8 genome assembly revealed the location of centromeres in the reference strain 70-15 of *M. oryzae*. Repeats, RNA-seq reads, and bisulfite sequencing reads were also mapped and are represented here for the comparative analyses. **(B)** A map showing seven chromosomes of *M. oryzae* with centromere locations marked on each chromosome. The chromosome length along with centromere length obtained from the ChIP-seq analysis is plotted to the scale on the available chromosome-wide 70-15 genome assembly. However, telomeres are shown as 10 kb regions on either side for each chromosome for visualization purpose.

Next, we analyzed the recently published PacBio genome assembly of the *M. oryzae* field isolate FJ81278 [42] to identify the centromere sequences and compare them with the 70-15 assembly. Mapping of CenpA ChIP-seq reads revealed nine distinct peaks in the FJ81278 genome assembly (S7 Fig and Table 1). Three of these enriched regions were present at the end of three separate contigs (Contig 3, 14 and 16). By comparing genome assemblies of 70-15 and FJ81278, we concluded that contigs 3 and 14 are most likely parts of the same chromosome and the CenpA-enriched regions observed in these two contigs represent a single centromere (*CEN4*). Synteny analysis also revealed that the CenpA peaks in Contig 11 and 16 belong to the same chromosome. However, Contig11 of FJ81278 assembly seems to be mis-assembled, since a part of this contig does not show synteny with any region of the 70-15 genome. Thus, we excluded this centromere (*CEN7*) region from further analysis.

We further compared the centromeres and the flanking regions from the genome assemblies of Guy11, 70-15, and FJ81278. Detailed synteny analyses revealed that the centromere flanking regions are conserved among these three isolates indicating that the overall position of centromeres is likely conserved in different strains/field isolates of *M. oryzae* (Fig 5A). However, a major part of the centromere sequences was absent in the 70-15 genome assembly as compared to Guy11 and FJ81278. It is important to note that the MG8 version of the 70-15 genome assembly is not complete and harbors a number of gaps. We believe that some of the centromere sequences are part of the unassembled Supercontig8.8 and are the CenpA-enriched regions observed in this fragment. The centromere sequences of Guy11 and FJ81278 isolates shared a high level of conservation with certain rearrangements. To explore this further, we performed a pair-wise comparison using sequences of respective centromeres from Guy11 and FJ81278 genomes. This analysis revealed that while most of the AT-rich sequence remains conserved between the two isolates, the repeat content varies significantly and accounts for almost all the observed rearrangements (Fig 5B and S2 Dataset). These results suggest that repeat elements might shape the structure of centromeres in different isolates even though they may not be an integral part of centromeres.

**Fig 5.**
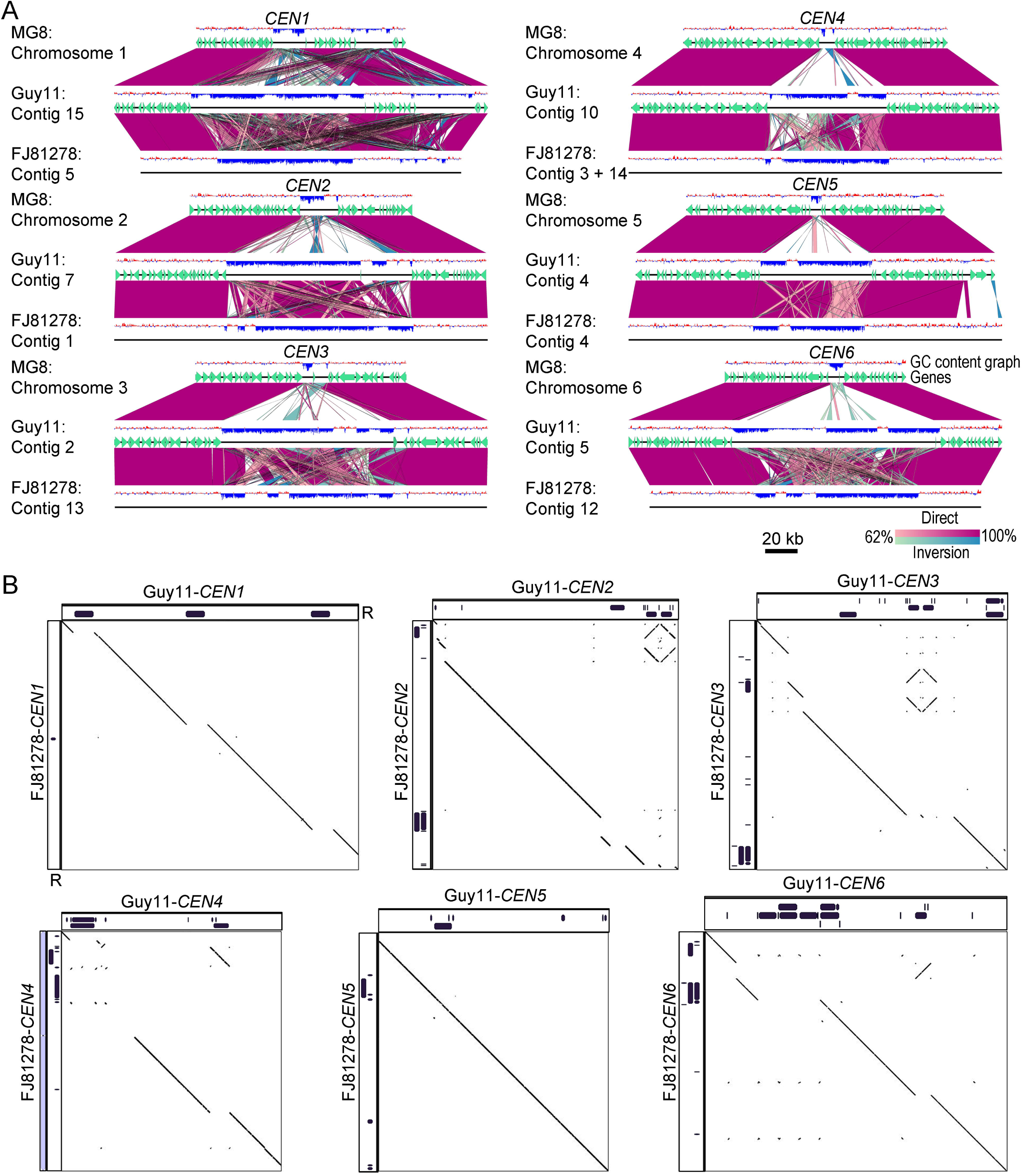
Centromere DNA sequences in *M. oryzae* isolates are similar, but vary in repeat content. **(A)** Synteny analysis across centromeres and their flanking regions revealed the conservation of centromere flanking regions indicating that the centromere location is maintained in different isolates of *M. oryzae*. The gene annotations for the FJ81278 assembly are not available and hence are not represented in the maps. This analysis also revealed that centromere sequences are largely excluded from the current MG8 genome assembly as compared to that of Guy11 or FJ81278. A 200 kb region (with respect to the Guy11 genome assembly) for each centromere is represented in the maps. A few centromere flanking genes were also found missing from Chr7 in the MG8 assembly. BLAST analysis revealed the presence of these genes in the unassembled supercontigs8.8 of the genome assembly. **(B)** Dot-plot analysis of respective centromeres revealed that centromere sequences share considerable similarities in Guy11 and FJ81278. As shown in the graphs, the breaks observed in the dot-plot analysis overlapped with the presence/absence of repeat elements. The complete sequence of *CEN4* in FJ81278 was generated by fusing the two fragments, one each from contig 3 and 14. The individual fragments are shown using grey bars and are separated by a small thin black bar (equal to 100 bp). R denotes the repeat panels for both Guy11 and FJ81278.

### Intra-and Inter-species comparison of *CEN* sequences in *Magnaporthales*

The analysis in different isolates of *M. oryzae* further validated that centromeres in this species comprise of long AT-rich and transcription-poor regions. Using these parameters, we decided to predict the centromeres in the wheat blast isolate B71 (Triticum pathotype of *M. oryzae;* MoT) as well as in *M. poae*, a root infecting pathogen that belongs to the *Magnaporthaceae* family [37]. Wheat blast isolate B71 genome was assembled to a chromosome level and exhibits a few chromosomal rearrangements as compared to the rice blast 70-15 genome assembly [48]. We identified seven putative centromeres, one in each chromosome, in the B71 genome. These centromeres in the wheat blast genome were long AT-rich regions (Fig 6A and Table 1). We then analyzed the centromere flanking regions between two genomes and found that centromere locations between the rice blast and wheat blast strains are conserved (Fig 6B). Further analysis revealed that the centromeres in wheat blast B71 also harbor a few repeats, but are defined primarily by AT-richness, similar to those observed in the rice blast isolates. We analyzed Scaffold 1 representing the mini-chromosome in B71 [48] and identified a putative centromere based on a long AT-rich region (40,795 – 109,519 bp) (Fig 6C, Table 1).

**Fig 6.**
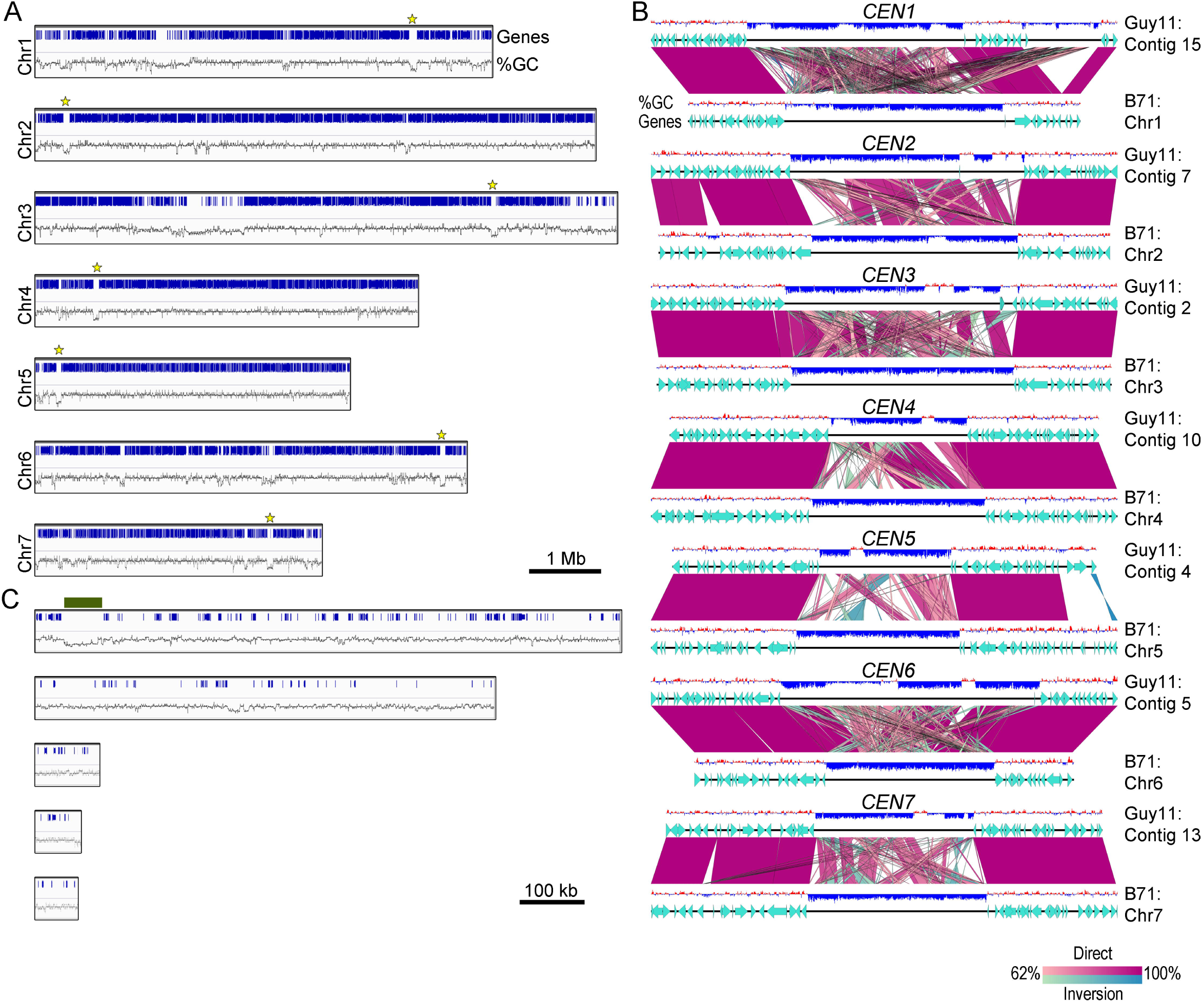
Predicted centromeres in the wheat blast genome. (A) Chromosome maps showing the location of centromeres in the genome of the B71 isolate of wheat blast. The AT-rich, gene-free centromere regions are marked by a yellow star in each chromosome. **(B)** Synteny analysis for centromeres and their flanking regions between the Guy11 and B71 genomes showed conservation of centromere locations therein. A 200-kb region is shown with respect to the Guy11 genome assembly. **(C)** Scaffolds representing the mini-chromosome in B71 were analyzed and are depicted with their gene and AT-richness graphs. A long AT-rich region (marked with dark green bar) in scaffold 1 (40,795 – 109,519) was identified and represents the putative centromere in the mini-chromosome in wheat blast.

Next, we extended our *in-silico* analysis to *M. poae* a distinct species within the *Magnaporthales* [33] and identified eight putative centromere regions across its genome (S8 Fig and Table 1). Three of these eight putative *CEN* regions are present at the end of different contigs. Since the chromosome number in *M. poae* is not established, it is uncertain whether all of these AT-rich regions represent bona fide centromeres in *M. poae*. We also found that these putative centromeres in *M. poae* harbor more repetitive DNA sequences than in *M. oryzae* even though the genomic repeat content of *M. poae* is only 1.1% as compared to 10.1% in *M. oryzae*. Unlike different isolates of *M. oryzae* that share a high level of centromere sequence conservation, the centromere sequences in *M. oryzae* and *M. poae* are highly divergent. Based on these results, we conclude that centromere DNA sequences in the *Magnaporthales* are rapidly evolving, whereas the properties of centromeric chromatin are likely conserved between the two species.

## Discussion

Blast diseases caused by *M. oryzae* is exceedingly disastrous not only to rice production worldwide but also to wheat and other graminaceous crops [35, 38]. Despite being such a vital plant pathogen, the fundamental cellular process of chromosome segregation is not well understood in this organism. In this work, we attempted to study the chromosome segregation machinery in *M. oryzae* at the molecular level. We tagged two evolutionarily conserved key kinetochore proteins in *M. oryzae* and studied their dynamics during different phases of the cell cycle at various developmental stages. We further identified the genomic loci that act as centromeres in this filamentous fungus. Based on the comparison of centromeric sequences among different host-adapted lineages of *M. oryzae,* and in the related species (*M. poae*), centromeres appear to be rapidly evolving in the *Magnaporthe* species complex, similar to those reported in several fungal genera [6, 8, 10, 12, 49, 50].

Kinetochores cluster together in a single locus at the nuclear periphery in many fungi. This locus is often referred to as the CENP-A-rich zone or CENP-A cloud [51, 52]. It has been proposed that such a nuclear subdomain with a high concentration of CENP-A favor centromere seeding on the chromosomal regions in close proximity to it, in the absence of a centromere-specific DNA sequence. In most budding yeasts, kinetochores are clustered throughout the cell cycle except in *C. neoformans,* which shows clustered kinetochores only during mitosis [19]. The kinetochore dynamics in *M. oryzae* is found to be similar to the “fission” yeast rather than that of the budding yeast species. It is possible that mitotic declustering of kinetochores is a feature of all yeasts/fungi that divide by septum formation. However, a more detailed analysis of kinetochore behavior in filamentous fungi like *N. crassa* and *Z. tritici* will be useful to establish this link. It is noteworthy that *Z. tritici* does not have a single centromere cluster, but kinetochores are arranged in multiple chromocenters, a process observed in some plant species [26, 53]. We also observed that kinetochores align along the mitotic spindle in *M. oryzae*, though a proper metaphase plate formation was not evident. A similar kinetochore arrangement was also observed in a basidiomycete *C. neoformans* [19]. The presence of such a structure in two evolutionarily distant fungal species suggests the existence of a transient formation of a structure, an arrangement alternative to the metaphase plate, across the fungal kingdom. In addition, co-localization of kinetochore proteins and SPBs revealed a close association between the two as observed in *S. pombe* [25]. Our results also suggest that a direct interaction between the SPBs and kinetochores may facilitate kinetochore clustering. The SPB-kinetochore interaction has been explored in other fungi and led to the identification of several uncharacterized proteins [22, 54-57]. It remains to be seen whether or not such interactions occur in *M. oryzae* as well.

Centromere DNA sequences, despite being associated with a conserved and essential function, are highly divergent across species [3]. The centromeres identified in *M. oryzae* further add to this diversity of centromere sequences. Our results show that centromeres in *M. oryzae* are long and AT-rich similar to those reported in *N. crassa* except that the centromeres are shorter in *M. oryzae* (57-109 kb) compared to *N. crassa* (150-300 kb) [12, 13]. The DNA methylation pattern observed in *M. oryzae* is similar to that of *N. crassa* as it is present at multiple loci in both the organisms and thus differs from that of *C. neoformans* where DNA methylation is restricted to only centromeres and telomeres [8, 13]. Additionally, a specific pattern of centromeric histone binding has been reported in *N. crassa,* but no such pattern exists in *M. oryzae*. Since centromere DNA sequences are generally repeat-rich, they are poorly assembled which restricts finer analysis of *CEN* DNA sequence. For example, centromeres in *Fusarium graminearum* are proposed to be AT-rich, similar to that of *M. oryzae* and *N. crassa* [12]. However, the exact nature of the centromere sequence of these regions remains unknown in *F. graminearum* due to sequence gaps in the genome assembly. Similarly, most of the centromere sequences are absent in the currently available 70-15 genome assembly of *M. oryzae*. Taken together, an improved genome assembly with complete chromosome-level sequence information is required for a better understanding of a complex genomic locus like the centromere.

Apart from filamentous fungi, AT-rich centromeres are present in other fungal species like *Malassezia sympodialis*, albeit the length of these regions is significantly smaller as compared to *M. oryzae* centromeres [32]. The CDEII element of point centromeres present in the budding yeast, *S. cerevisiae*, is also highly AT-rich [58]. A recent study reports the presence of AT-rich centromeres of varying lengths in diatoms [30]. Furthermore, the 171-bp alpha satellite repeat DNA present in human centromeres is also AT-rich in nature [59]. Overall, these results suggest that AT-richness favors centromere function in many organisms. Intriguingly, *in vitro* experiments suggest that CENP-A binds with a lower affinity to an AT-rich DNA sequence [60]. In contrast, the same study also revealed that the CENP-A chaperone Scm3 has a higher affinity towards AT-rich sequences. With more AT-rich centromeres being characterized, identifying the exact role of AT-rich sequences in centromere function is critical.

Regional centromeres of many organisms, including *M. oryzae*, do not share any common DNA sequence motifs. Rather, non-DNA sequence determinants mark centromeres in an epigenetic manner in many organisms. Some epigenetic determinants of centromere identity in fungi include early replicating regions of the genome [61-63], proximity to DNA replication origins [64], DNA replication initiator proteins [65], homologous recombination-repair proteins [64, 66] and proteins that facilitate kinetochore clustering by tethering kinetochores to SPBs [22, 54]. Factors that favor local folding and looping of chromatin may also add to the process of centromere specification [4, 52, 67]. Repeats and transposons have been shown to play an essential role in centromere evolution [68-70]. Previous reports in *M. oryzae* suggested the presence of multiple clusters of repeat elements across the genome [43, 47]. These studies also proposed that repeats play an important role in *M. oryzae* genome evolution and its association with the host. In this study, we find that the centromere location is close to these repeat clusters in some but not all chromosomes. Our results raise the possibility that centromere sequences in *M. oryzae* are prone to repeat-mediated evolution.

A comparison between two *M. oryzae* isolates, Guy11 and FJ81278, revealed that while the overall *CEN* DNA sequence between the two rice blast isolates is very similar, the repeat content at the centromeres of orthologous chromosomes varies widely. It is known that centromere DNA sequence among isolates of *N. crassa* can be different [12]. The *CEN* sequences identified here would pave the way for a more detailed comparative analysis of centromeres in diverse isolates of *M. oryzae.* Such analyses will provide valuable insights into centromere evolution in this species and the potential impact of host factors on this process. A comparative genome analysis between *M. oryzae* and *M. poae* revealed the presence of a higher density of repeats in the predicted *CEN* regions in the latter. Overall, these results suggest that while the centromere DNA sequence properties, not the DNA sequence *per se,* remain conserved in the fungal order *Magnaporthales*, the centromere architecture is divergent and might have been shaped by the repeat elements. Further studies will provide more insights into the evolution of centromere DNA sequences and its possible link to host adaptation and variability in virulence within this important family of cereal killers.

## Materials and Methods

### Fungal strains and culture conditions

Wild-type *M. oryzae* strain Guy11 (MAT1-2; a kind gift from Lebrun group, France) was used as the parent strain for all the experiments conducted in this study (except for the results shown in S3 Fig, S5 and S6 Movie that were performed using B157 strain). All strains were propagated on prune-agar (PA) medium or complete medium (CM) as described previously [71]. For conidiation, fungal strains were grown on PA agar plates at 28 °C for two days in the dark followed by exposure to continuous light at room temperature for five days. Conidia were harvested using an inoculation loop by gently scraping the culture surface with sterile distilled water. The resulting conidial suspension was filtered through two layers of Miracloth (Calbiochem) to remove mycelial debris and was adjusted to obtain the required concentration after counting with a hemocytometer. *Agrobacterium tumefaciens*-mediated transformation (ATMT) of *M. oryzae* was carried out as described previously [71, 72]. Transformants were screened for antibiotic resistance using the respective selection media, i.e., CM with 250 μg/ml Hygromycin B, Basal media with 50 μg/ml ammonium glufosinate (selection for Basta) or Chlorimuron-ethyl (selection for Sulfonylurea). Transformants were verified for correct genomic integration by diagnostic PCR and sequencing. The strains thus validated and used in this study are listed in S2 Table. The plasmids and primers used for epifluorescence labeling in *M. oryzae* strains are listed in S3 and S4 Table, respectively. The detailed information about the plasmid construction is available in the S1 Text.

### Microscopy and image processing

Unless otherwise stated, live cell microscopy imaging was performed on a motorized inverted Nikon Eclipse Ti-E microscope with perfect focus system equipped with a Yokogawa CUS-X1 spinning disk confocal system and a CFI Plan Apo VC 100x/1.4 NA oil objective lens. The images were captured using 16-bit digital Orca-Flash4.0 sCMOS camera (Hamamatsu Photonic K.K) and at 491 nm 100 mW (for green fluorescence) and at 561 nm 50 mW (for red fluorescence) laser illumination operated by MetaMorph Premier Software (Ver. 7.7.5, Universal Imaging). The maximum projection images were obtained from Z stacks of 0.5 µm-spaced sections using the built-in MetaMorph module. Image processing was performed using Imaris (Bitplane) and Fiji (https://imagej.net/Fiji). A Live-SR module (Gataca Systems), which is based on optically demodulated structured illumination technique with online processing, was additionally mounted on the same spinning disk confocal system during image acquisitions for Figure 2A. The high-resolution images for Figure 2B and 2C were acquired using an Andor Dragonfly high-speed confocal microscope equipped with an iXon888 EMCCD camera and a 100x oil objective lens. The raw images were immediately processed using the integrated Fusion software and the in-built deconvolution feature.

### Chromatin immunoprecipitation

The ChIP experiment was performed using the protocol described previously with a few modifications [73]. *M. oryzae* strain expressing GFP-CenpA or CenpC-GFP fusion protein was grown in 150 ml complete media for three days at 28 °C with continuous shaking at 150 rpm. Fungal mycelia were collected using two layers of Miracloth (Calbiochem), and the harvested mycelia were washed with sterile water. Mycelia were cross-linked by suspending them in 1% formaldehyde solution in 20 mM HEPES/pH7.4 for 20 min with continuous shaking at 100 rpm. Glycine was added to the suspension at a final concentration of 0.125M, and the mix was further incubated at room temperature for an additional 10 min. Cross-linked mycelia were harvested using Miracloth and rinsed with water. The excess water was removed by gently patting the mycelia mass with paper towels followed by snap-freezing in liquid nitrogen. The frozen mass was then stored at −80°C until used. For each ChIP experiment, 80-100 mg of frozen mycelia were ground in liquid nitrogen using mortar-pestle, and powdered mycelia were resuspended in 1 ml of nuclei isolation buffer (10 mM MES-KOH/10 mM NaCl/10 mM KCl/2.5 mM EDTA pH8.0/250 mM sucrose/0.1 mM spermine/0.5 mM spermidine-free base/1 mM DTT). Nuclei were separated from debris by filtering them through two layers of Miracloth and pelleted by centrifugation at 13000 rpm for 10 min at 4 °C. The pellet was resuspended in 1 ml of lysis buffer (50 mM HEPES pH 7.5/150 mM NaCl/1 mM EDTA/0.1% Na-deoxycholate/1% Triton-X and 0.1% SDS). The resuspended nuclei were subjected to sonication using a Bioruptor (Diagenode) for 60 cycles of the 30s on and 30s off bursts at the high level, and the fragmented chromatin (300 - 600 bp) was isolated by centrifugation at 13000 rpm, 10 mins at 4 °C. A part of the chromatin fraction (100 μl) was used for input DNA (I) preparation and the remaining chromatin solution was divided into two halves of 450 μl each. In one of the tubes, 20 μl of GFP-TRAP beads (ChromoTek) were added for immunoprecipitation (+) while the other tube was incubated with 20 μl of blocked agarose beads (ChromoTek) to use as the negative control (-). The tubes were incubated at 4 °C for 12 h on a rotator. The beads were then washed, and bound chromatin was eluted in 500 μl of elution buffer (1% SDS, 0.1M NaHCO_3_). All three fractions (I, + and -), were de-crosslinked and DNA was isolated using phenol: chloroform extraction followed by ethanol precipitation. The precipitated DNA was air dried and dissolved in 25 μl of MilliQ water containing 25 μg/ml RNase (Sigma-Aldrich). I and + samples were subjected to ChIP-sequencing for GFP-CenpA ChIP in *M. oryzae*. CenpC-GFP ChIP samples (I, + and -) were subjected to qPCR with centromere-specific primers along with a non-centromeric primer set. The fold enrichment for CenpC at the centromere was plotted using GraphPad Prism software.

### Analysis of sequencing data

GFP-CenpA ChIP sequencing was performed at the Clevergene Biocorp. Pvt. Ltd., Bengaluru, India. A total of 52,604,910 and 38,186,320 150 bp paired-end reads were obtained for Input and IP samples, respectively. The reads were mapped to Guy11 PacBio genome (GCA_002368485.1), MG8 genome assembly (GCA_000002495.2) and FJ81278 genome assembly (GCA_002368475.1) using Geneious 9.0 (http://www.geneious.com/) with default conditions. Each read was allowed to map only once randomly anywhere in the genome. The alignments were exported to bam files, sorted and visualized using Integrative Genomics Viewer (IGV, Broad Institute). The images from IGV were imported into Adobe Photoshop (Adobe systems) and scaled for representation purpose. The ChIP-sequencing reads have been deposited under NCBI BioProject Accession ID PRJNA504461.

RNA-sequencing data (SRR1568068) from a previous study was downloaded from the NCBI website and was aligned to genomes using RNA-seq alignment default parameters in Geneious 9.0. Similarly, bisulfite sequencing data (SRR653493) was obtained from NCBI, and the reads were aligned to genomes using Geneious default aligner. The aligned files were exported into bam files and visualized using IGV. The RNA-seq reads were plotted in logarithmic scale for the visualization purpose. GC-content was calculated using Geneious 9.0 with a sliding window size of 200 bp. The data was exported as wig files and further visualized using IGV.

For *M. poae*, the GC content and RNA-seq data were plotted as described for *M. oryzae*. The RNA-seq data was downloaded from NCBI (SRR057701) and was aligned to the reference genome (GCA_000193285.1).

The repeats annotations presented in this study were done based on repeat sequences described in a previous study [37]. BLASTn analysis was carried out using these repeat sequences, results were sorted, and hits with 100% query coverage were extracted. The hits were mapped onto the respective genome assemblies of Guy11, FJ81278, B71 or *M. poae* and visualized using IGV.

### Synteny analysis

The synteny analysis between centromere flanking regions was performed using Easyfig software [74] (http://mjsull.github.io/Easyfig/). The graphs were plotted with default condition (except the color settings), and GC percentage was plotted as 200 bp sliding window. The dot-plot analysis for centromere sequences was carried out using the gepard software with a 100-bp window [75].

## Acknowledgments

We thank Clevergene Biocorp Pvt. Ltd., Bengaluru for generating the CenpA ChIP-sequencing reads. We thank the Fungal Pathobiology group at TLL for useful discussions and suggestions; and the TLL Bioimaging Facility for help with confocal microscopy and image analyses.

