## Supplementary Information for "Cellular Dynamics and Genomic Identity of Centromeres in the Cereal Blast Fungus"

**Supplementary Methods**

**Plasmid construction for epifluorescent-tagged strains**

**Histone H1-mCherry (nuclear marker):** The PCR amplified mCherry open reading frame (ORF), without the start codon, was cloned into the plasmid vector pFGL822 (Addgene #58225), which contains the Basta (Phosphinothricin/Glufosinate) resistance cassette, using KpnI and BamHI restriction enzymes. Next, histone H1 ORF (without the stop codon) and its 3’UTR sequence were PCR amplified with the corresponding primers (listed in the Table S4) and ligated sequentially to generate the final plasmid pFGL1170R (Addgene #116896).

**CenpC-GFP and CenpC-mCherry (kinetochore markers):** The GFP ORF (without the start codon) was introduced into plasmid pFGL821 (Addgene #58223), which contains the Hygromycin B resistance gene. The C-terminal part of CenpC ORF (without the stop codon) and 1 kb of 3’ UTR were subsequently cloned into the plasmid to obtain CenpC-GFP reporter plasmid pFGL1079 (Addgene #116897). CenpC-mCherry construct pFGL1169R (Addgene #116899) was generated in a similar fashion to that of hH1-mCherry by cloning the CenpC ORF and the 3’ UTR.

**GFP-CenpA (kinetochore marker):** A codon-optimized Tet-off regulator cassette was commercially synthesized and cloned into the backbone plasmid pFGL1252 (Addgene #118991). Subsequently, a hygromycin resistance cassette was ligated in pFGL1252 to obtain the Tet-off plasmid pFGL1252_TetOFF (Hyg) (Addgene #118992). The GFP ORF (without the stop codon) was introduced immediately after the modified Tet-off cassette to obtain pFGL1252_TetGFP (Hyg) (Addgene #118993). The entire CenpA ORF (lacking the start codon) along with its 3’UTR and 1 kb of 5’ homology arm was cloned and ligated sequentially into pFGL1252_TetGFP(Hyg) to generate the Tet-off controlled GFP-CenpA expressing plasmid pFGL1258 (Addgene #116898).

**GFP-TubA and mCherry-TubA (microtubule marker):** The TubA ORF (without the ATG start codon) with its 3’UTR was PCR amplified and cloned immediately after the *GFP* in pFGL1259 (Addgene #118996). The promoter of TubA was then cloned upstream of the *GFP* coding sequence to get the final plasmid pFGL1260 (Addgene #116900). The mCherry-TubA construct pFGL1260R (Addgene #116901) was generated similarly by cloning the same fragments in pFGL1259R (Addgene #118995).

**Alp6-mCherry (Spindle pole body/MTOC marker):** A marker-fusion reporter construct, pFGL1269R (Addgene #118994), was used as the backbone vector for this plasmid construct. The C-terminal part of Alp6 ORF (without the stop codon) and 1 kb of its 3’ UTR were cloned into requisite sites upstream and downstream of the *mCherry* respectively, yielding the final plasmid pFGL1344 (Addgene #116902).

**Supplementary Figure Legends**

**S1 Fig. Identification of CenpA and CenpC in *M. oryzae*.** **(A)** A schematic representation of the kinetochore complex. For simplification, only the CenpA and CenpC are highlighted. The identified kinetochore proteins in *M. oryzae* are listed in Table S1. **(B)** Alignment of CenpA protein sequence from *M. oryzae* (Mo), with those of *Drosophila melanogaster* (Dm), *Mus musculus* (Mm), *Homo sapiens* (Hs), *Saccharomyces cerevisiae* (Sc), *Ustilago maydis* (Um), *Candida albicans* (Ca), *Cryptococcus neoformans* (Cn), *Schizosaccharomyces pombe* (Sp) and *Neurospora crassa* (Nc). The C-terminal region of CenpA is an evolutionarily conserved histone-fold domain (HFD), whereas the N-terminus exhibits a high level of sequence divergence. **(C)** Multiple sequence alignment of CenpC protein sequence from *M. oryzae* (Mo) with other species. CenpC protein sequences revealed conservation of the CenpC box and the DNA binding “Cupin” domain.

**S2 Fig. CenpA is essential for cell growth and viability in *M. oryzae*. (A)** Growth characteristics and colony morphology of the strain MGYF01 and the wild-type *M. oryzae* Guy11 in the absence (-Dox) or presence (+Dox) of doxycycline (5 mg/L). **(B)** Epifluorescence confocal imaging of the strain MGYF06 showing the organization of nuclei (H1-mCherry), kinetochores (GFP-CenpA) and microtubules (GFP-TubA). In interphase, the microtubule signals are localized in the cytoplasm, whereas a single clustered dot of GFP-CenpA co-localizes with chromatin marker, H1-mCherry in mycelia as well as conidia. **(C)** Epifluorescence confocal imaging of the strain MGYF09 showing the organization of SPB (Alp6-mCherry), kinetochores (GFP-CenpA) and microtubules (GFP-TubA). During mitosis, the SPB signals flank the mitotic spindle. The kinetochore (GFP-CenpA) signals cannot be distinguished from the initially strong signal of the spindle (GFP-TubA). However, as the mitotic spindle elongates, and the signal is diffused towards the end of mitosis, the GFP-CenpA signals co-localize with those of Alp6-mCherry at the poles. Also, see movie S4. The images shown are maximum projections of 0.5 µm-spaced *z* stacks. Scale bars = 5 μm.

**S3 Fig. Subcellular localization and dynamics of CenpA and CenpC during pathogenic development in *M. oryzae*.** **(A)** Time-lapse images showing a mitosis event during appressorium formation in *M. oryzae* isolate B157. *M. oryzae* conidia were incubated on the hydrophobic coverslip to allow appressorium development. The mitotic division was recorded after 4 hpi, and images captured at 20-second intervals. Also, see Movie S5. **(B)** Time-lapse images showing the occurrence of mitosis during *in planta* growth of *M. oryzae* isolate B157. *M. oryzae* conidia were incubated on rice sheath, and the images were acquired at 44 hpi at 13-second intervals. Also, see Movie S6. The epifluorescence images shown here are maximum projections from Z-stacks consisting of 0.5 μm-spaced planes. Scale bars = 5 μm.

**S4 Fig. Identification of centromeres in the wild-type *M. oryzae* strain Guy11*.*** GFP-CenpA ChIP-seq revealed the identity of centromeres in *M. oryzae*. Shown here is the CenpA binding pattern across contigs longer than 100 kb in Guy11 genome assembly. Contigs 1-17 (>500 kb in length) are shown with a different scale whereas contigs 18-28 (<500 kb, ≥100 kb in length) are represented with a different scale for visualization purpose. The presence of repeat elements, GC-content, level of transcription, and DNA methylation across all the contigs are also plotted.

**S5 Fig. Dot-plot analysis of *M. oryzae* (Guy11) centromeres.** Self-dot-plot analysis for each of the seven centromere regions was performed and plotted. Both *x*- and *y*-axis represent the length of each centromere.

**S6 Fig. *M. oryzae* strain 70-15 genome is syntenic to Guy11 genome. (A)** Chromosome-wide synteny block map of the reference MG8 (strain 70-15) genome assembly with contigs from the Guy11 genome assembly. **(B)** Alignment of CenpA ChIP-sequence reads to the unassembled region (supercontigs 8.8) from the chromosome-wide MG8 genome assembly identified two significantly enriched regions.

**S7 Fig. CenpA ChIP-seq read mapping identified centromere locations in *M. oryzae* isolate FJ81278.** Graphs showing the enrichment of CenpA in FJ81278 genome assembly. The enriched regions overlapped with AT-rich regions. The locations of CenpA enriched centromeres are marked. The graphs are plotted with two different scales for the visualization purpose.

**S8 Fig. Centromere identification in *Magnaporthe poae*.** Based on the RNA-seq reads, AT-richness and repeat content data, centromeres were identified (marked) in *M. poae* genome, and the graphs showing the same are plotted for the *M. poae* genome assembly. Contigs longer than 500 kb are only represented here.

**S1 Fig**

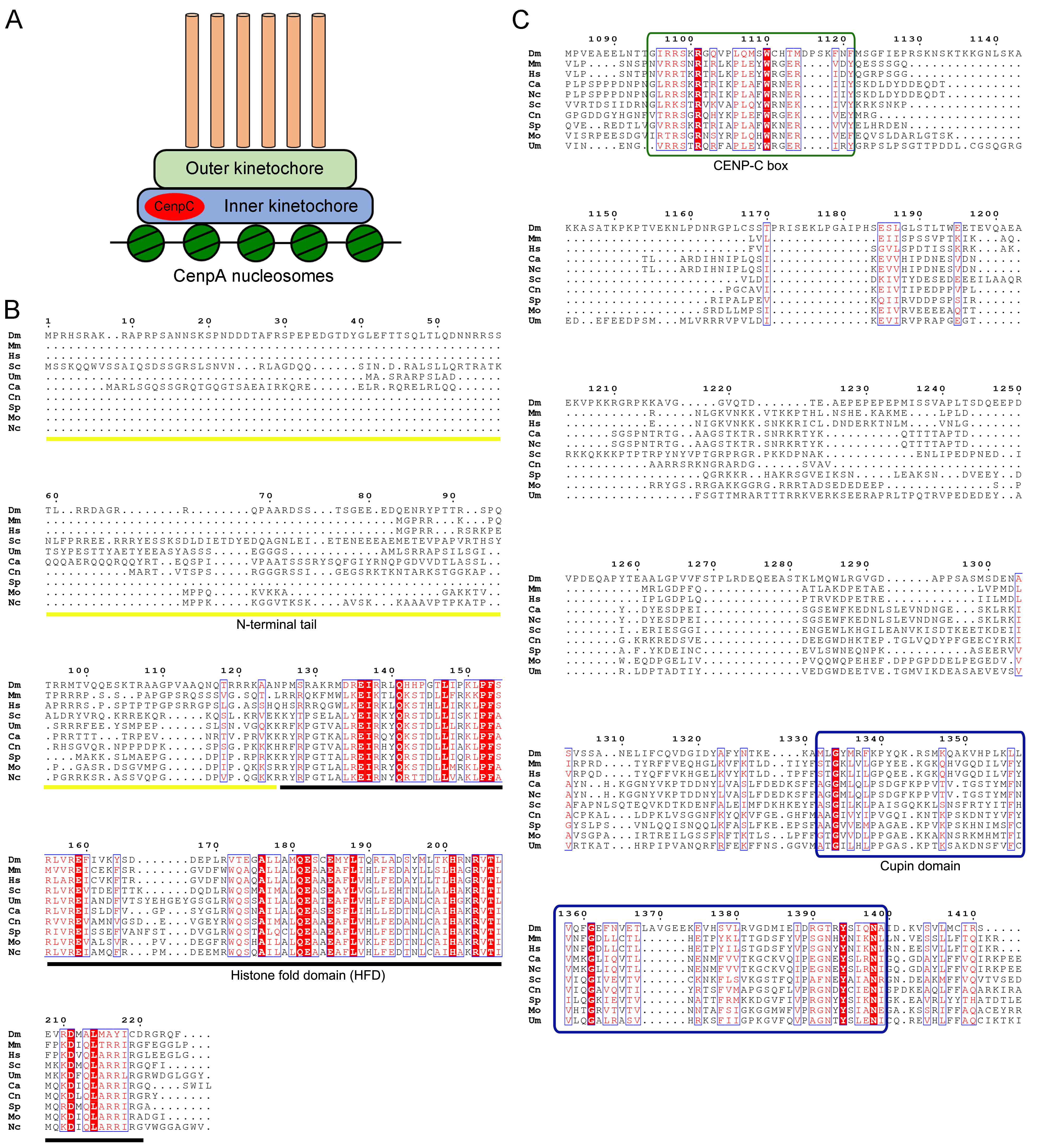

**S2 Fig**

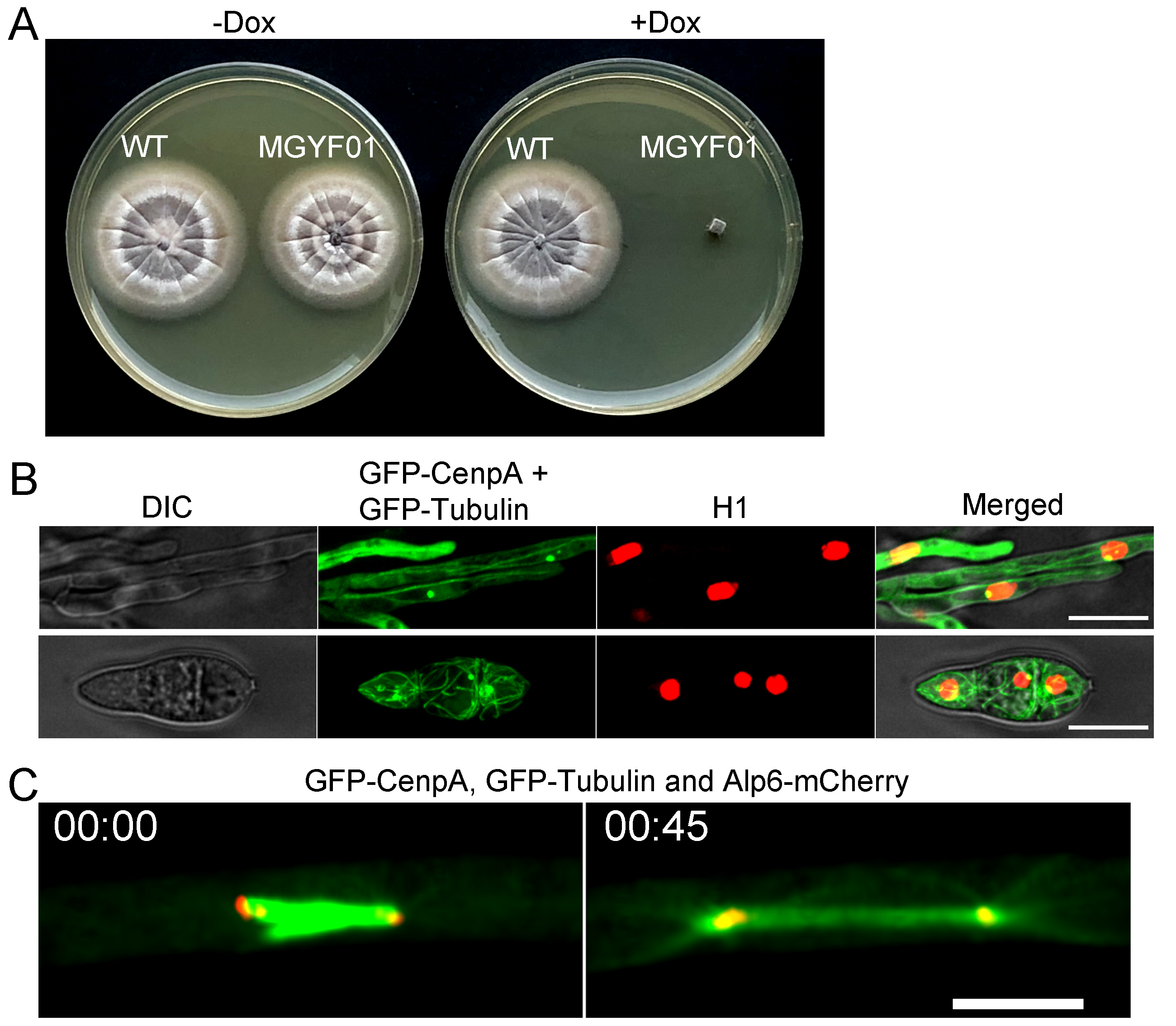

**S3 Fig**

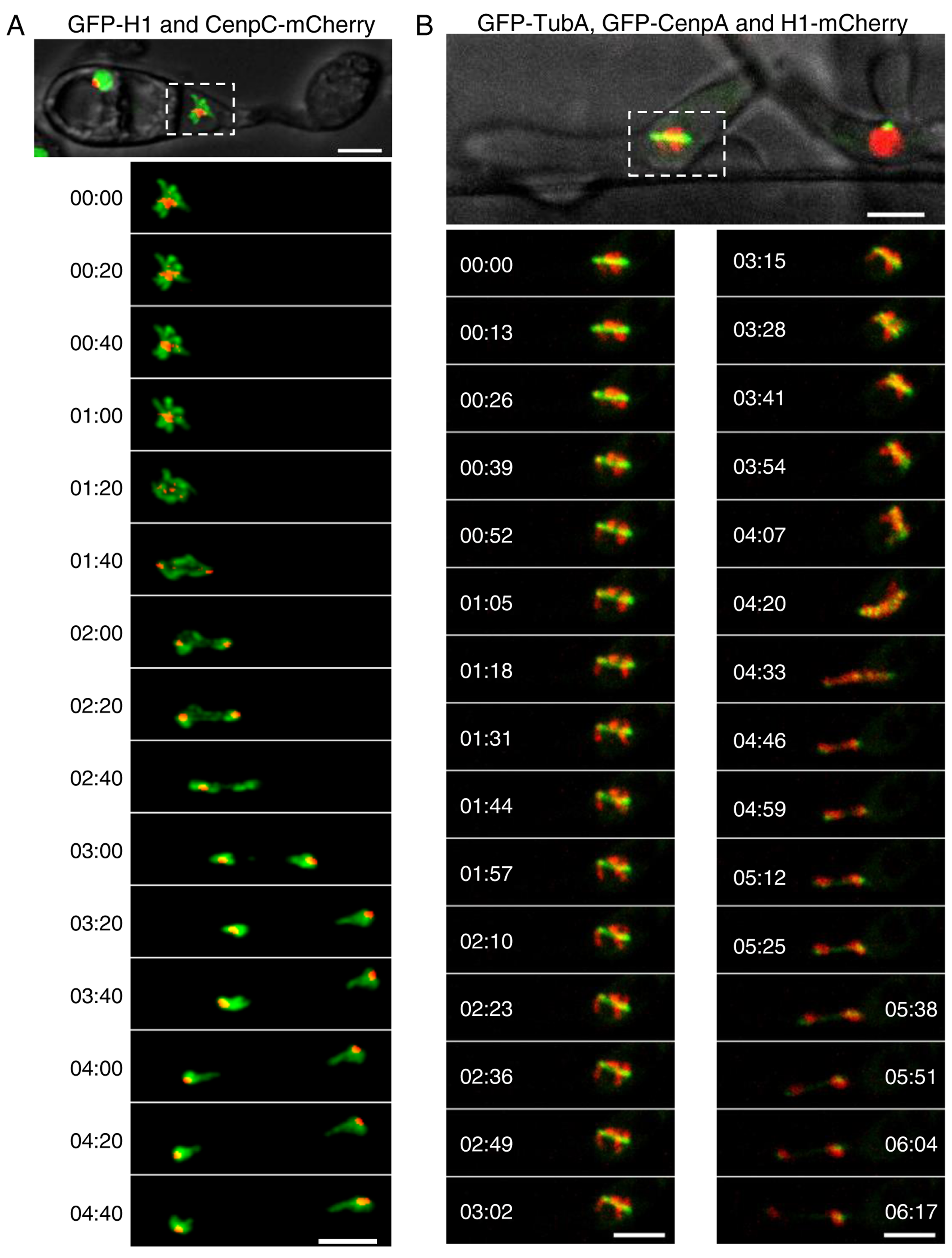

**S4 Fig**

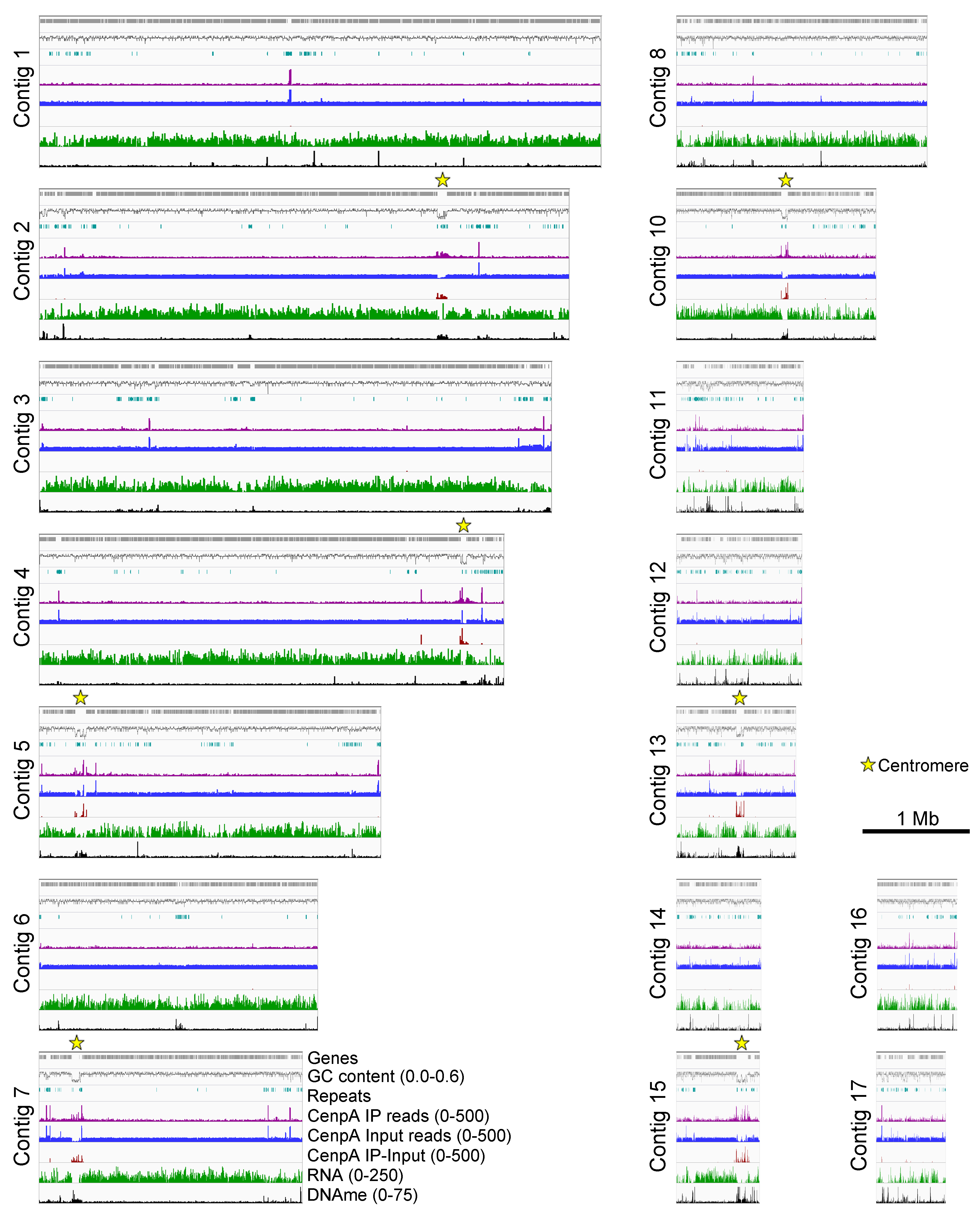

Continued next page…

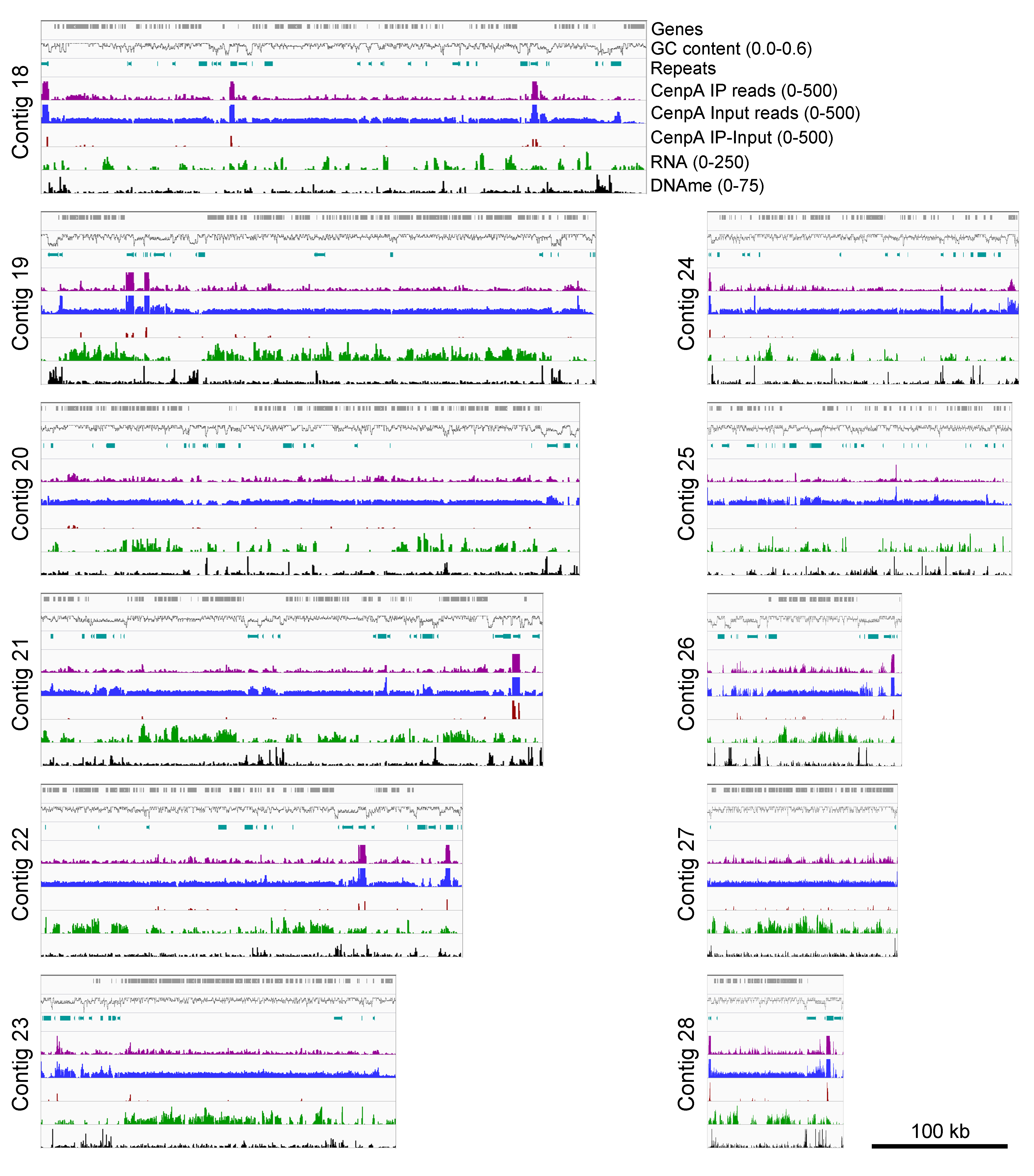

**S5 Fig**

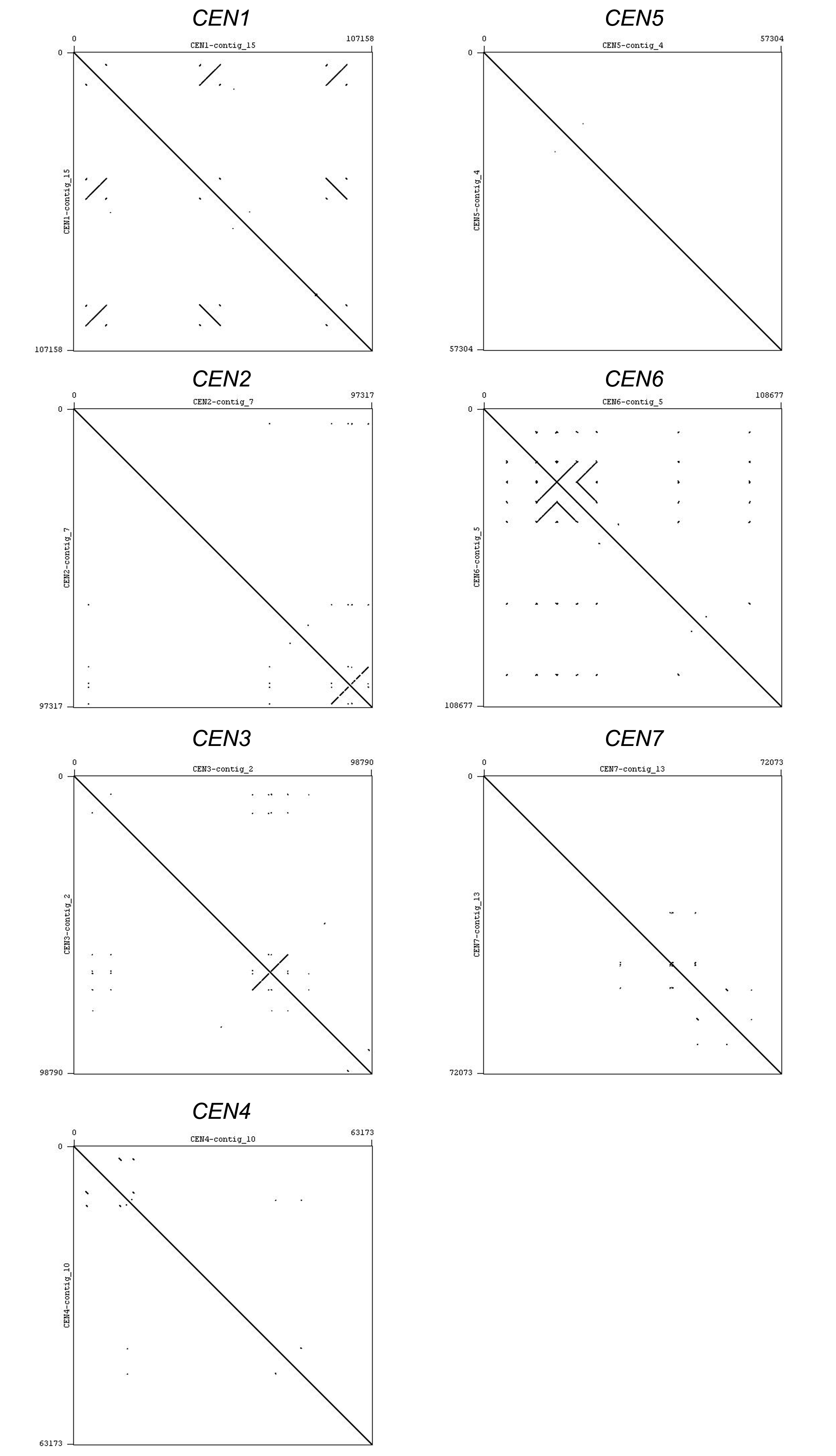

**S6 Fig**

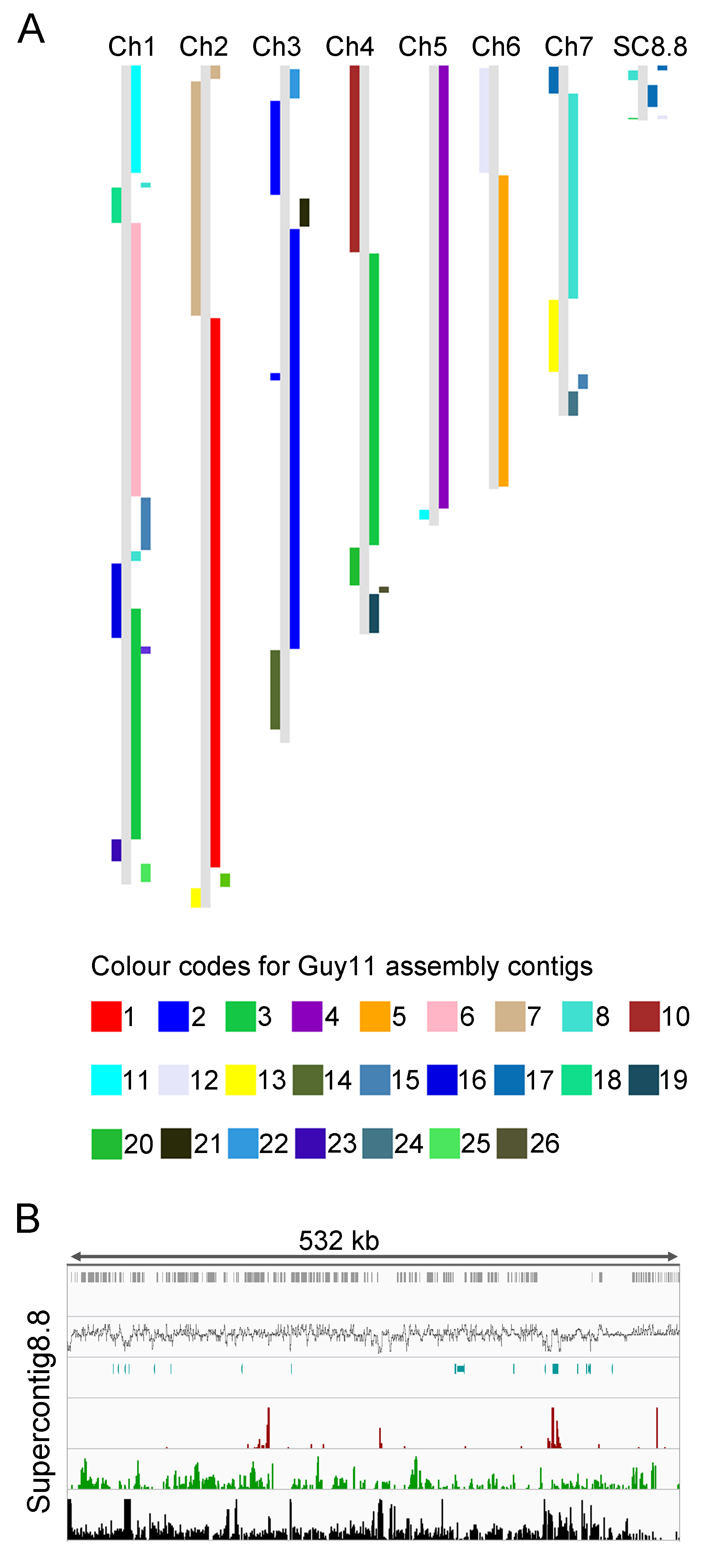

**S7 Fig**

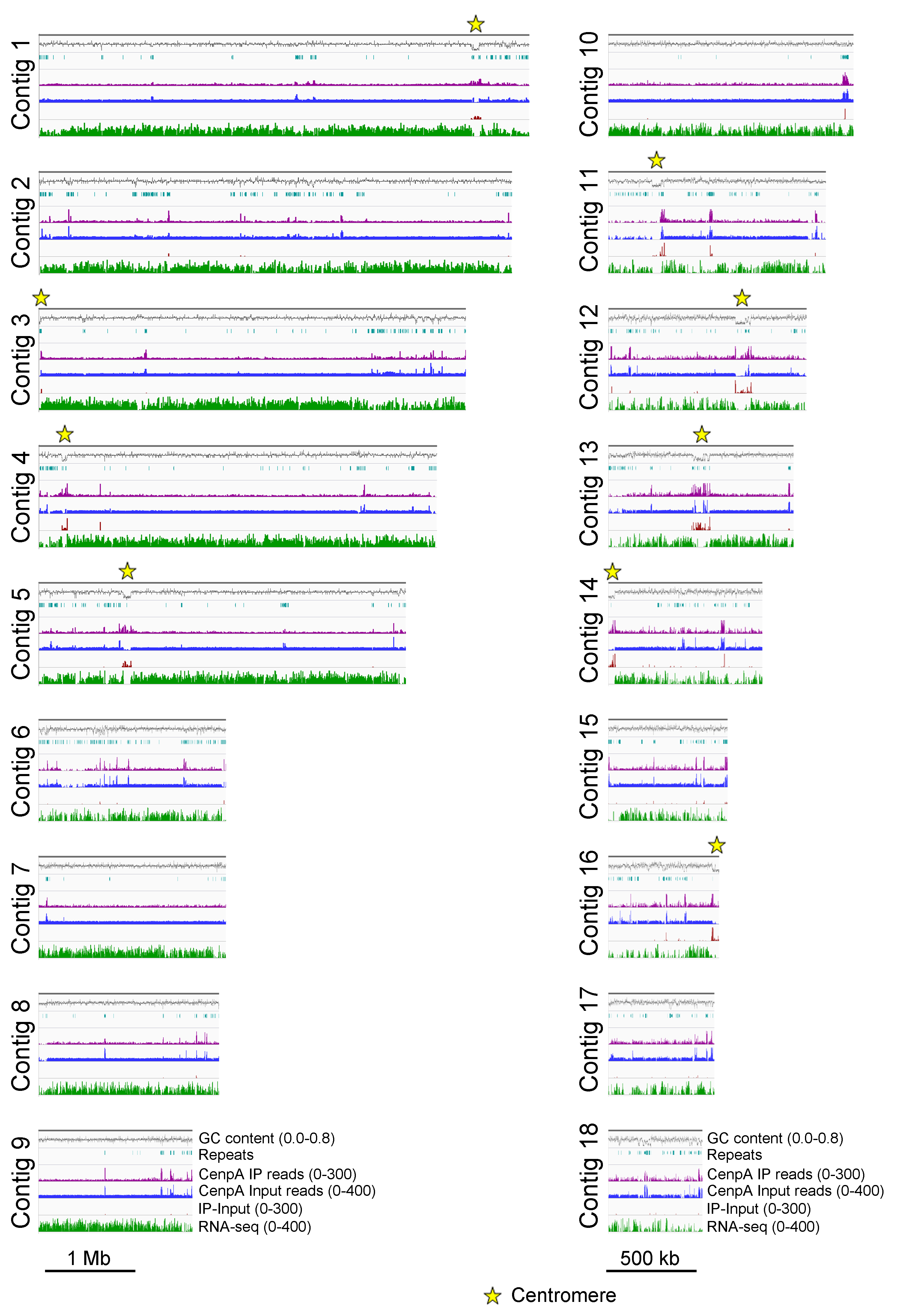

**S8 Fig**

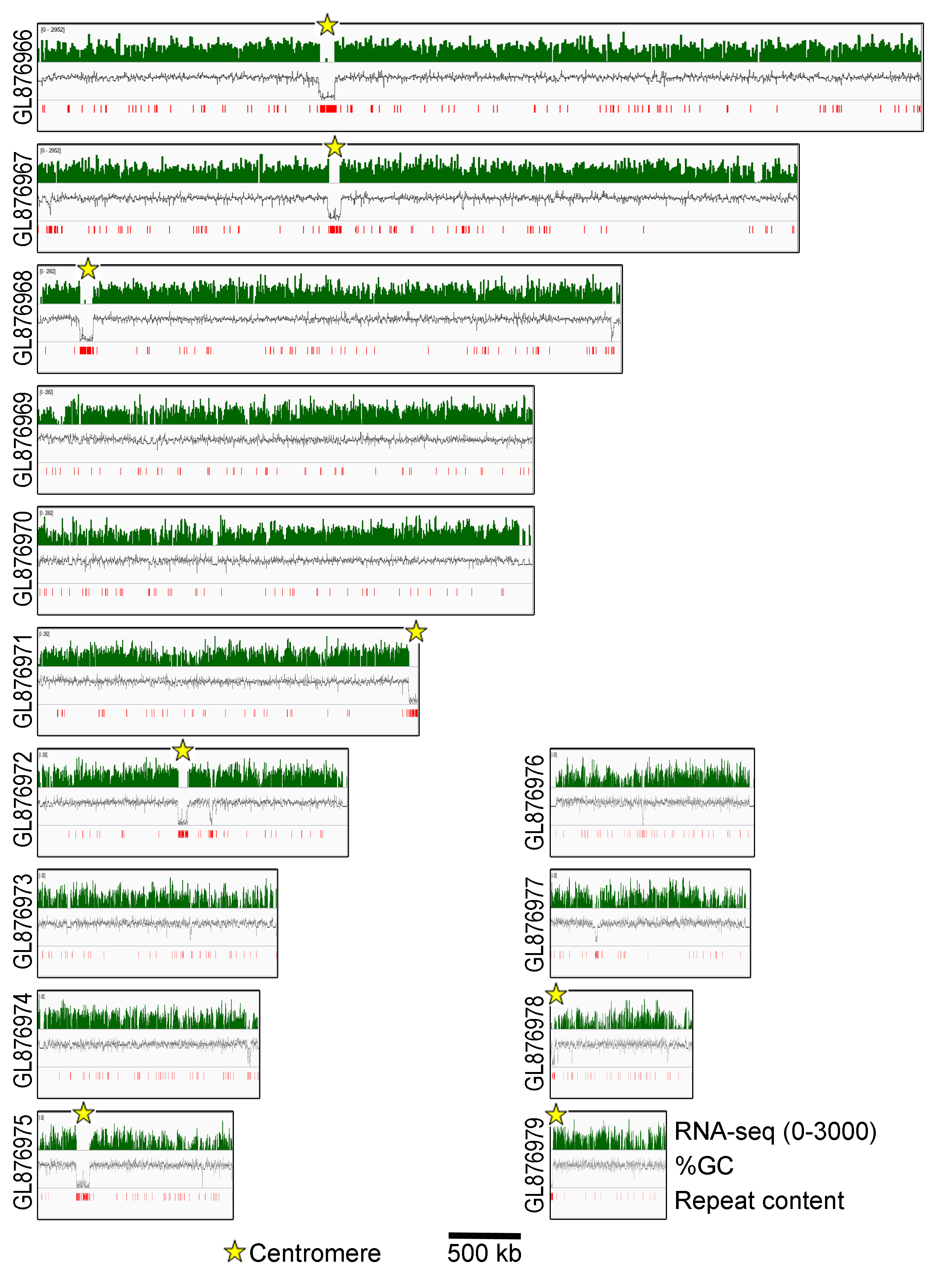

**Supplementary Movies**

**S1 Movie.** Dynamics of kinetochores (GFP-CenpA) and microtubules (mCherry-TubA) during mitosis in *M. oryzae*.

**S2 Movie.** Kinetochores (GFP-CenpA) undergoing unclustering during mitosis in *M. oryzae*.

**S3 Movie.** Kinetochore dynamics (GFP-CenpA) showing the segregation of sister kinetochores and reclustering during the mitotic division in *M. oryzae*.

**S4 Movie.** Dynamics of kinetochores (GFP-CenpA), microtubules (GFP-TubA) and SPBs (Alp6-mCherry) during mitosis in *M. oryzae*.

**S5 Movie.** Kinetochore (CenpC-GFP) and chromatin (mCherry-histone H1) dynamics during mitosis in appressorium of *M. oryzae* isolate B157.

**S6 Movie.** Dynamics of kinetochore (GFP-CenpA), microtubules (GFP-TubA) and chromatin (H1-mCherry) during *in planta* growth of B157 strain.

**Supplementary Datasets**

**S1 Dataset. Analysis of repeats/repetitive elements in *M. oryzae* (Guy11) and *M. poae* genome assemblies.**

**S2 Dataset. Percent identity comparison of centromere sequences from Guy11, FJ81278 and B71.**

**S1 Table. Kinetochore proteins in *M. oryzae*.**

| **Kinetochore protein/protein complex** | ***S. cerevisiae*** | ***H. sapiens*** | ***M. oryzae*** |
| --- | --- | --- | --- |
| **Dam1 Complex**  Dad1  Dad2  Dad3  Dad4  Dam1  Duo1  Spc19  Spc34  Ask1  Hsk3 | + | **Ska complex** | MGG_12092  MGG_02522  MGG_06996  MGG_16761  MGG_00874  MGG_17765  MGG_09127  MGG_00887  MGG_07143  MGG_15008 |
| **KNL1/Spc105** | + | + | MGG_03693 |
| **Ndc80 Complex**  Ndc80  Nuf2  Spc24  Spc25 | + | + | MGG_01027  MGG_01848  MGG_06076  MGG_08566 |
| **Mtw1 Complex**  Mtw1  Dsn1  Nnf1  Nsl1 | + | + | MGG_06304  MGG_08211  MGG_04669  MGG_00906 |
| **CENP-O/P/Q/U**  CENP-O  CENP-P  CENP-Q  CENP-R  CENP-U | + | + | MGG_12605  -  MGG_06713  -  MGG_15811 |
| **CENP-H/I/K complex**  CENP-H  CENP-I  CENP-K | + | + | MGG_04487  MGG_09521  MGG_17440 |
| **CENP-T**  **CENP-X**  **CENP-W**  **CENP-S** | + | + | MGG_02732  MGG_16148  MGG_11869  MGG_15064 |
| **CENP-C** | + | + | MGG_06960 |
| **CENP-A** | + | + | MGG_06445 |

‘+’ present; ‘-’ absent

**S2 Table. List of strains used in this study.**

| **Strain** | **Brief description** | **Purpose** | **Selection** |
| --- | --- | --- | --- |
| Guy11 | Wild type (MAT1-2) | Parent stain for fungal transformation | None |
| MGYF01 | Guy11 transformed with pFGL1258 | GFP-CenpA tagging under Tet-off promoter regulation | Hyg |
| MGYF02 | Guy11 transformed with pFGL1079 | CenpC-GFP tagging | Hyg |
| MGYF03 | MGYF01 transformed with pFGL1170R | GFP-CenpA and H1-mCherry | Hyg, Bar |
| MGYF04 | MGYF02 transformed with pFGL1170R | CenpC-GFP and H1-mCherry | Hyg, Bar |
| MGYF05 | MGYF01 transformed with pFGL1169R | GFP-CenpA and CenpC-mCherry | Hyg, Bar |
| MGYF06 | MGYF03 transformed with pFGL1260 | GFP-CenpA, H1-mCherry and GFP-TubA | Hyg, Bar, Sur |
| MGYF07 | MGYF01 transformed with pFGL1260R | GFP-CenpA and mCherry-TubA | Hyg, Sur |
| MGYF08 | MGYF01 transformed with pFGL1344 | GFP-CenpA and Alp6-mCherry | Hyg, Bar |
| MGYF09 | MGYF08 transformed with pFGL1260 | GFP-CenpA, Alp6-mCherry and GFP-TubA | Hyg, Bar, Sur |

**S3 Table. Plasmid constructs used in this study.**

| **Constructs**  **(Addgene #)** | **Targeted gene** | **Description/purpose** | **Selection marker** |
| --- | --- | --- | --- |
| pFGL1170R  (116896) | hH1 (MGG_12797) | C-terminal tagging with mCherry at the native locus, as a nuclear marker. | Basta |
| pFG1079  (116897) | CenpC (MGG_06960) | C-terminal tagging with GFP at the native locus, as a kinetochore marker. | Hygromycin |
| pFGL1258  (116898) | CenpA (MGG_06445) | N-terminal tagging with GFP with a Tet-OFF cassette at the native locus, as a kinetochore marker. | Hygromycin |
| pFGL1169R  (116899) | CenpC (MGG_06960) | C-terminal tagging with mCherry at the native locus, as a kinetochore marker. | Basta |
| pFGL1260  (116900) | TubA (MGG_11412) | N-terminal tagging with GFP at the ILV2 locus, as a microtubule marker. | Sulfonylurea |
| pFGL1260R  (116901) | TubA (MGG_11412) | N-terminus tagging with mCherry at the ILV2 locus, as microtubule marker. | Sulfonylurea |
| pFGL1344  (116902) | Alp6 (MGG_01815) | C-terminal tagging with mCherry at its native locus, as an MTOC marker. | Basta |

**S4 Table. Oligonucleotide primers used in this study.**

| **Primer Name** | **Sequence (5' to 3')** | **Application** |
| --- | --- | --- |
| GFP_F4_KpnI | CATC**GGTACC**GTGAGCAAGGGCGAGGAGCTGT | GFP without start codon |
| GFP_R720_BamHI | CGC**GGATCC**TTACTTGTACAGCTCGTCCATGC |  |
| GFP_R717_BamHI | CGC**GGATCC**CTTGTACAGCTCGTCCATGCC | GFP without stop codon |
| mCherry_F4_KpnI | CTC**GGTACC**GTGAGCAAGGGCGAGGAGGATAA | mCherry without start codon |
| mCherry_R711_BamHI | CGC**GGATCC**TTACTTGTACAGCTCGTCCATGC |  |
| mCherry_R708_BamHI | CGC**GGATCC**CTTGTACAGCTCGTCCATGCCGC | mCherry without stop codon |
| hH1_F4_XhoI | CGT**CTCGAG**CCTCCCAAGAAGGAAACC | 5’ homologous arm of hH1 (ORF without stop codon) |
| hH1_R1039_EcoRI | GTC**GAATTC**TGCGGCGGGTGCCTCGGC |  |
| hH1_endF_PstI | CCG**CTGCAG**TAAAGGGACGCTGACGAACTT | 3’ homologous arm of hH1 |
| hH1_R+871_HindIII | CAT**AAGCTT**CTTTCTTTGACGGGAAAGGGA |  |
| CenpA_F(-804)_EcoRI | GGT**GAATTC**AGTCGCAGGTACATCTCATTA | 5’ homologous arm of CenpA |
| CenpA_R0_EcoRI | GTG**GAATTC**TTTATGTCGGTTTCTATATGGTTTC |  |
| CenpA_F4_BamHI | ATA**GGATCC**CCACCACAAAAAGTAAAGAAGG | 3’ homologous arm of CenpA (ORF+3’ UTR) |
| CenpA_R+425_XbaI | GTT**TCTAGA**AAAGCAGTCCCCAGAGTAACTT |  |
| CenpC_F1704_EcoRI | CCG**GAATTC**GAGGACGAAGAACC | 5’ homologous arm of CenpC (without stop codon) |
| CenpC_R2253_KpnI | GAT**GGTACC**GCTGCTTTCAGTCATTTCGTCC |  |
| CenpC_F2254_PstI | CTG**CTGCAG**TAATTCATCGCGGTGGGTTGGTC | 3’ homologous arm of CenpC |
| CenpC_R+966_HindIII | TGT**AAGCTT**CTGGCCCTCCCTCATTAT |  |
| TubA_F(-1042)_XhoI | GCA**CTCGAG**GCCGCCGGTGTAATTCATGGTGACT | TubA promoter |
| TubA_R3_KpnI | CTC**GGTACC**CATTGTGGATTCTAGGCACTTTTCTCAG |  |
| TubA_F4_BamHI | TCC**GGATCC**AAAGGCGAGGTAGGTGATGCTT | TubA (ORF+3’ UTR) |
| TubA_R+559_XbaI | GCGC**TCTAGA**GGGTGAAACCAAGTATGT |  |
| Alp6_F1876_EcoRI | CTC**GAATTC**GCAATATGCAAGCCCAGAAGTGC | 5’ homologous arm of Alp6 (without stop codon) |
| Alp6_R2876_KpnI | AAC**GGTACC**GTCTCCAGCACGCTCGCCCCTG |  |
| Alp6_endF_XbaI | TGC**TCTAGA**TAAGTAGTTTGAGAAGGAATGACGC | 3’ homologous arm of Alp6 |
| Alp6_R+967_HindIII | ATG**AAGCTT**GGGTCGATGAGTGGTTGGGTT |  |
| HR_F 15.1 | CGCCGTATATTTTCTTCCG | *CEN1* ChIP qPCR |
| HR_R 15.1 | CGAAAATTTCGAAATTCTGCC |  |
| HR_F 7.2 | GACCAAACCTCCTATTACC | *CEN2* ChIP qPCR |
| HR_R 7.2 | GTAAAGGGGAATATTGCCG |  |
| HR_F 2.2 | TGCGCTAAAGATTCCGGG | *CEN3* ChIP qPCR |
| HR_R 2.2 | TCGTTGGTTATATTAAGCGG |  |
| HR_F 10.1 | GGCACCGGAAACCTTTTC | *CEN4* ChIP qPCR |
| HR_R 10.1 | ACGTTGTACGCAATTGGAC |  |
| HR_F 4.1 | CGCACAAAGATTATAAGGTAG | *CEN5* ChIP qPCR |
| HR_R 4.1 | CTCAAATACGTTATTTTAAACAG |  |
| HR_F 5.1 | TTTAACGTCGATTACCATTTC | *CEN6* ChIP qPCR |
| HR_R 5.1 | TAAGACAAATGGGTTCAAATC |  |
| HR_F 13.1 | CGATAAAAACGCGTTTGCG | *CEN7* ChIP qPCR |
| HR_R 13.1 | GAATTTCGTCGGGGTTTTC |  |
| Mg.CEN6.ctrl.FP | CTTTTGCTCAAGAAGCAGCG | Non-*CEN* ChIP qPCR |
| Mg.CEN6.ctrl.RP | TCGGTGTGCGTTTCGTGCTG |  |
| HR_F_62 | GAGTTTATCTCCCGGCTTG | MGG_01054 ChIP qPCR |
| HR_R_63 | CAGAGTACGGATCGCCTC |  |

Note: The restriction enzyme sites are underlined.
